# An empirical pipeline for personalized diagnosis of Lafora disease mutations

**DOI:** 10.1101/2021.03.26.437206

**Authors:** M. Kathryn Brewer, Maria Machio-Castello, Rosa Viana, Jeremiah L. Wayne, Andrea Kuchtová, Zoe R. Simmons, Sarah Sternbach, Sheng Li, Maria Adelaida Garcia-Gimeno, Jose Serratosa, Pascual Sanz, Craig W. Vander Kooi, Matthew S. Gentry

**Affiliations:** Department of Molecular and Cellular Biochemistry, University of Kentucky College of Medicine, Lexington, KY 40536, USA; Lafora Epilepsy Cure Initiative, Epilepsy and Brain Metabolism Center, and Center for Structural Biology, University of Kentucky College of Medicine, Lexington, KY 40536, USA; Institute for Research in Biomedicine (IRB Barcelona), 08028 Barcelona, Spain; Neurology Laboratory and Epilepsy Unit, Department of Neurology, IIS Fundación Jiménez Díaz, Autónoma University, and Centro de Investigación Biomédica en Red de Enfermedades Raras (CIBERER), Madrid 28040, Spain; Instituto de Biomedicina de Valencia, CSIC and Centro de Investigación Biomédica en Red de Enfermedades Raras (CIBERER), Valencia 46010, Spain; Department of Medicine, University of California at San Diego, La Jolla, CA 92093, USA; Department of Biotechnology, Escuela Técnica Superior de Ingeniería Agronómica y del Medio Natural (ETSIAMN), Universitat Politècnica de València, 46022 Valencia, Spain

**Keywords:** laforin, Lafora disease, epilepsy, phosphatase, glycogen storage disease, carbohydrate-binding module, personalized medicine, glycogen

## Abstract

Lafora disease (LD) is a fatal, insidious metabolic disorder characterized by progressive myoclonic epilepsy manifesting in the teenage years, rapid neurological decline, and death typically within ten years of onset. Mutations in either *EPM2A*, encoding the glycogen phosphatase laforin, or *EPM2B*, encoding the E3 ligase malin, cause LD. Whole exome sequencing has revealed many *EPM2A* variants associated with late-onset or slower disease progression. We established an empirical pipeline for characterizing laforin missense mutations *in vitro* using complimentary biochemical approaches. Analysis of 26 mutations revealed distinct functional classes associated with different outcomes supported by multiple clinical cases. For example, F321C and G279C mutations have attenuated functional defects and are associated with slow progression. This pipeline allows rapid characterization and classification of novel *EPM2A* mutations, enabling clinicians and researchers to rapidly utilize genetic information to guide treatment of LD patients.

## INTRODUCTION

Lafora disease (LD) is a fatal progressive myoclonic epilepsy. LD patients develop normally until their adolescent years, when generalized seizures and myoclonic jerks begin. Over time, patients experience increasingly severe and frequent epileptic episodes, cognitive decline, ataxia and aphasia, leading to childhood dementia, and a vegetative state ^1^. Antiseizure drugs are only palliative, and most patients do not live beyond age 30. A hallmark of LD is the presence of polyglucosan bodies in most tissues, known as Lafora bodies (LBs) ^2^. Work by multiple laboratories has demonstrated that LBs are the etiological agent driving LD ^3–6^. Thus, LD is classified as a member of the larger glycogen storage disease (GSD) family, which affects 1 in 20,000-43,000 newborns ^7^.

LD patients carry mutations in either the *Epilepsy progressive myoclonus 2A (EPM2A)* gene encoding laforin or the *EPM2B* gene encoding malin. Laforin is the founding member of the glucan phosphatase family and dephosphorylates glycogen, a soluble glucose-storage molecule synthesized by eukaryotic cells ^8–10^. We previously defined the structural basis for laforin glycogen binding and dephosphorylation and characterized the quaternary structure of laforin and its dynamics ^11^. Laforin also directly interacts with and is ubiquitinated by malin, an E3 ubiquitin ligase ^12,13^. It has been suggested that laforin also serves as a central glycogen-associated scaffolding protein, interacting with multiple partner proteins important for glycogen metabolism ^14^ and that a possible cause of LD is the lack of a functional laforin-malin complex ^15^.

A significant amount of genetic information has emerged from whole exome sequencing of rare diseases ^16^. Over one hundred distinct LD-causing mutations in *EPM2A* have been identified including missense and nonsense mutations and insertions/deletions (indels) (http://projects.tcag.ca/lafora/) ^17,18^. While the number of reported mutations grows yearly, many mutations have only been identified in a single patient, often with compound heterozygosity, and published clinical details can be sparse ^19,20^. This lack of data makes genotype-phenotype correlations difficult. Computational algorithms can be employed to predict pathogenicity of a variant; however, these algorithms range in performance, and results from different programs often do not correlate ^21–23^. In many cases, experimental strategies can define patient-specific pathogeneses, explain differences in disease severity, and enable personalized medicine ^24^. This approach has been successful in the Cystic fibrosis (CF) field. Effects of disease-causing genetic variants were difficult to predict using *in silico* tools, and were instead carefully defined through many basic biological and biochemical studies ^25–29^. Now, CF mutations are classified based on their functional effect(s), and mutation-specific therapies are used to treat patients ^25^.

LD was previously described as a largely homogenous disease course irrespective of the patient’s mutation ^18,30^. Before the genetic loci were identified, if patients presented with progressive myoclonic epilepsy and lived beyond the age of 30, then a diagnosis of LD was ruled out ^31^. Even now, adult patients with milder progressive myoclonus epilepsy typically do not undergo LD screening ^32^. However, late-onset and slowly progressing LD have now been confirmed in older patients by genetic testing ^31,33^. With the increasing use of genetic testing to confirm LD diagnosis, only recently has the neurology community begun to explore the heterogeneous progression of LD patients ^32,34^.

In this report, we describe four additional LD patients with *EPM2A* missense mutations displaying this newly recognized spectrum of disease severity. Three patients presenting with a classic aggressive LD course and one displaying a protracted course. To define the molecular basis for these differences and others previously reported, we establish a pipeline to rapidly profile laforin functions and analyze 26 laforin missense mutations, including the four in this clinical report. The mutations segregate into five classes based on their biochemical effects and provide an explanation for classic and slowly progressing forms of LD. Moving forward, novel mutations can be quickly classified using our empirical pipeline and distinguished from benign polymorphisms. This system can be used to guide patient-specific clinical care and the deployment of therapeutics, which is needed to maximize the benefit from emerging novel LD therapeutics that are being developed ^34–39^.

## RESULTS

### Clinical features of four LD patients

We describe four patients from four families with *EPM2A* mutations that experienced varying clinical courses to highlight the heterogeneous progression observed in LD patients and introduce progression categories (Table 1). All four patients are compound heterozygous for a missense mutation and a nonsense mutation. Patient 1 (E28K/W165X) developed visual and generalized tonic-clonic seizures at the age of 10 years, with absence seizures manifesting at 11 years and myoclonic seizures at 13 years. He developed cognitive problems at 15 years and one year later started with speech difficulties and ataxia. He was wheelchair-bound at 17 years. He needs assistance for all activities of daily living and has no social interaction. Patient 2 (W32G/R241X) suffered his first visual seizure followed by a generalized tonic-clonic seizure at 10 years and absence seizures at 13 years. At age 14 years he developed myoclonic jerks, began suffering from cognitive impairment, and dropped out of school. When he was 16 years old, he presented gait unsteadiness and became wheelchair-bound. Presently he is bedridden, mute and has continuous myoclonus. Patient 3 (W32G/W165X) experienced cognitive problems at age 14 years and epilepsy with myoclonic and generalized tonic-clonic seizures at 16 years. When he was 17 years old, he became wheelchair-bound and speechless. Patient 4 (G279C/Q55X) suffered a first generalized tonic-clonic seizure at age 25 years, and after 7 years of evolution has no cognitive or motor impairment.

**Table 1.** Clinical features, genetic findings and classification of selected *EPM2A* patients.

| Patient | Age at onset | Nucleic acid change | Protein mutation | clinical state 5 years after disease onset |
| --- | --- | --- | --- | --- |
| 1 | 10 | c.82G>A<br>c.495G>A | E28K<br>W165X | Epilepsy and cognitive impairment |
| 2 | 10 | c.94T>G<br>c.721C>T | W32G<br>R241X | Epilepsy and cognitive impairment |
| 3 | 14 | c.94T>G<br>c.495G>A | W32G<br>W165X | Bedridden and mute |
| 4 | 25 | c.835G>A<br>c.163C>T | G279C<br>Q55X | Epilepsy with no cognitive or motor impairment |

All 4 patients were compound heterozygous for a missense mutation and a nonsense mutation. While patients 1 and 2 presented a classic progression and patient 3 an ultra-rapid developing subtype, patient 4 displayed a slower progression. Late-onset LD was also reported in a compound heterozygous patient with EPM2A mutations G279C/R241X ^40^. Therefore, it is possible that the G279C mutation is less detrimental to the function of laforin than the missense mutations carried by patients 1, 2 and 3, E28K and W32G.

### Pathogenicity predictions of *EPM2A* missense mutations

To define the dysfunctional basis for *EPM2A* missense mutations, 26 variants spanning the entire laforin protein were initially selected for analysis (Table 1 and Fig. 1a). Some variants are associated with a slower disease course (G279C and P211L) or a very late-onset phenotype (F321C). These mutations were initially analyzed using three *in silico* tools to predict mutation pathogenicity: PolyPhen-2 ^41^, SDM ^42^, and CUPSAT ^43^. PolyPhen-2 produced predictions for the mutants ranging from benign to probably damaging, with the majority predicted as deleterious substitutions (Table 2). Although the mutations F5S, V7A, F84L, N148Y, and E210K were all predicted with high confidence to be benign variants, all of these mutations are associate with typical LD^44–46^, with the exception of F5S for which no clinical information has been published (http://projects.tcag.ca/lafora/). SDM and CUPSAT predict changes in free energy (ΔΔG) induced by a missense mutation using different methodologies. Both programs estimated that the majority of mutations would be destabilizing (Table 2). However, a positive ΔΔG (increased stability) for W32G, Y294N, and P301L was predicted by at least one program, and we previously showed using purified proteins that all of these mutations were significantly destabilizing ^11^. Strikingly, there was no obvious clustering of mutations or consensus predictions between the three *in silico* tools (Fig. 1b).

**Fig. 1.**
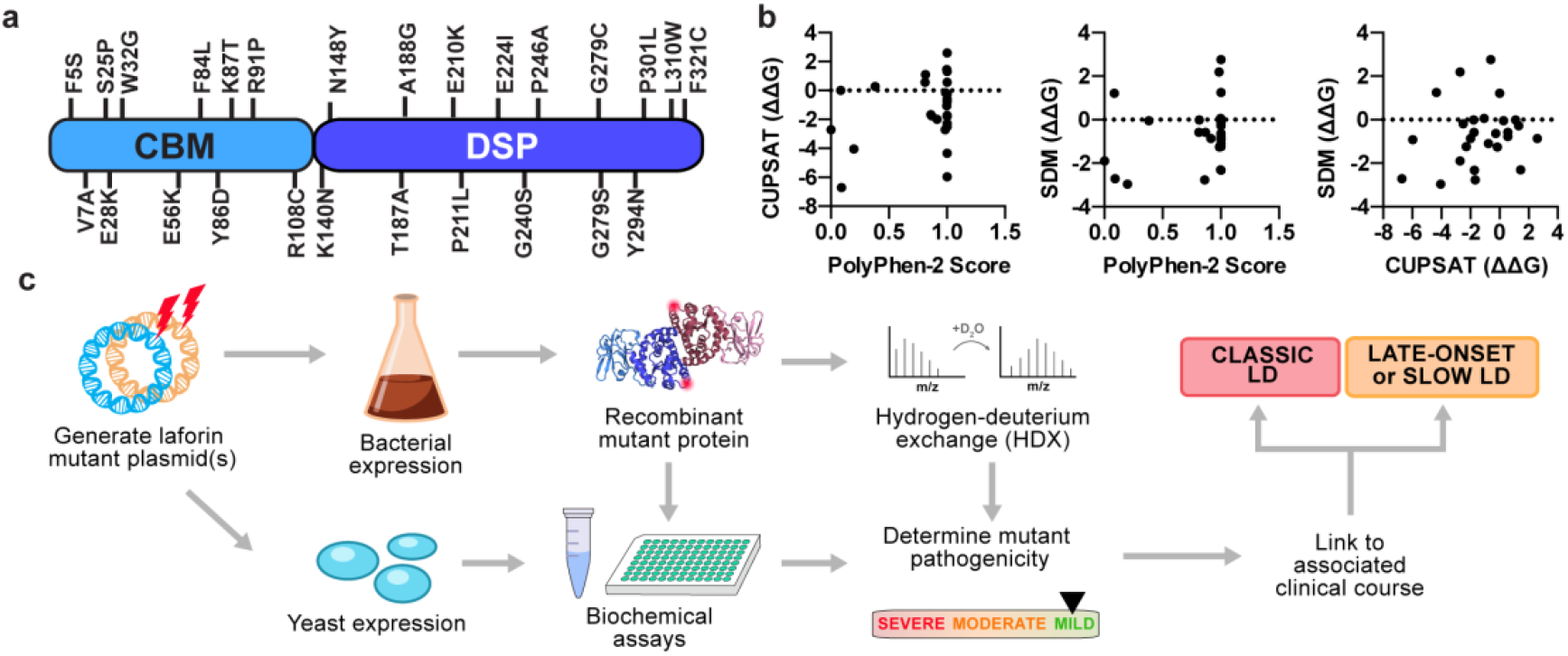
Pathogenicity predictions and a biochemical pipeline for *EPM2A* missense mutations. (a) Laforin is a bimodular protein with a CBM (carbohydrate binding module) and DSP (dual specificity phosphatase) domain. LD missense mutations selected for study are shown mapped to the primary sequence of laforin. (b) Correlation analysis between PolyPhen-2 pathogenicity score and ΔΔG (kcal/mol) predictions by CUPSAT and SDM for the 26 mutations selected for analysis (see Table 2). (c) Empirical pipeline for characterizing missense mutations *in vitro*. The mutations were introduced into bacterial and yeast expression plasmids. Bacterially purified recombinant protein was used for biochemical assays and hydrogen deuterium exchange (HDX) studies. Yeast two-hybrid assays were performed to study laforin interactions with binding partners. The biochemical profile of the mutant was then compiled to determine the severity of the mutation, which is then linked to the clinical course of the patient.

**Table 2.** Predicted pathogenicity of selected *EPM2A* missense mutations. Asterisk(s) indicates this mutation has been explicitly associated with a slow or late-onset phenotype, either in the compound heterozygous (*) or homozygous (**) state. The Lafora Epilepsy Mutation Database can be found here: http://projects.tcag.ca/lafora/. CUPSAT and SDM cutoff scores: stabilizing/neutral >=0 (green); destabilizing, between −2 and 0 (orange); destabilizing <−2 (red). For CUPSAT results, ΔΔG was predicted for residues in both subunits of the laforin structure (chain A and C or PDB:4rkk) and the values in the table represent an average.

|  |  | PolyPhen-2 |  | CUPSAT |  | SDM |  |
| --- | --- | --- | --- | --- | --- | --- | --- |
| | | Probability Score | Prediction | $\Delta\Delta G$ (kcal/mol) | Predicted overall stability | $\Delta\Delta G$ (kcal/mol) | Outcome |
| F5S | Lafora Epilepsy Mutation Database<sup>17</sup> | 0.198 | Benign | -4.045 | Destabilizing | -2.96 | Reduced stability |
| V7A | <sup>44</sup> | 0.092 | Benign | -6.71 | Destabilizing | -2.71 | Reduced stability |
| S25P | <sup>60</sup> | 1 | Probably damaging | 0.56 | Stabilizing | -0.78 | Reduced stability |
| E28K | <sup>30</sup><br>Current report | 1 | Probably damaging | -0.76 | Destabilizing | -1.09 | Reduced stability |
| W32G | <sup>30</sup><br>Current report | 1 | Probably damaging | -0.595 | Destabilizing | 2.76 | Increased stability |
| E56K | <sup>19</sup> | 1 | probably damaging | 2.605 | Stabilizing | -0.87 | Reduced stability |
| F84L | <sup>45</sup> | 0.003 | Benign | -2.705 | Destabilizing | -1.89 | Reduced stability |
| Y86D | <sup>40</sup> | 0.863 | possibly damaging | -1.68 | Destabilizing | -2.77 | Reduced stability |
| K87T | <sup>19</sup> | 1 | probably damaging | -0.26 | Destabilizing | -0.63 | Reduced stability |
| R91P | <sup>84</sup> | 0.988 | Probably damaging | -0.145 | Destabilizing | -1.26 | Reduced stability |
| R108C | <sup>30,60</sup> | 1 | Probably damaging | 1.31 | Stabilizing | -0.28 | Reduced stability |
| K140N | <sup>46</sup> | 0.915 | Possibly damaging | -1.975 | Destabilizing | -0.86 | Reduced stability |
| N148Y | <sup>46</sup> | 0.085 | Benign | 0 | No effect | 1.21 | Increased stability |
| T187A | <sup>61</sup> | 0.999 | Probably damaging | -1.06 | Destabilizing | 0.06 | Increased stability |
| A188G | <sup>18</sup> | 1 | Probably damaging | -5.965 | Destabilizing | -0.92 | Reduced stability |
| E210K | <sup>46</sup> | 0.381 | Benign | 0.28 | Stabilizing | -0.04 | Reduced stability |
| P211L* | Lafora Epilepsy Mutation Database<sup>17</sup> | 0.867 | Possibly damaging | -1.74 | Destabilizing | -0.58 | Reduced stability |
| E224I | <sup>60</sup> | 0.812 | Possibly damaging | 0.585 | Stabilizing | -0.57 | Reduced stability |
| G240S | <sup>45</sup> | 0.815 | Possibly damaging | 1.11 | Stabilizing | -0.01 | Reduced stability |
| P246A | <sup>18</sup> | 0.985 | Probably damaging | -2.71 | Destabilizing | 2.19 | Increased stability |
| G279C* | <sup>40</sup><br>Current report | 1 | Probably damaging | -1.76 | Destabilizing | -0.01 | Reduced stability |
| G279S* | <sup>30,45,85,86</sup> | 1 | Probably damaging | -1.73 | Destabilizing | -2.32 | Reduced stability |
| Y294N | <sup>45,85</sup> | 0.999 | Probably damaging | 1.44 | Stabilizing | -2.3 | Reduced stability |
| P301L | <sup>45</sup> | 1 | Probably damaging | -4.335 | Destabilizing | 1.26 | Increased stability |
| L310W | <sup>46</sup> | 1 | Probably damaging | -2.275 | Destabilizing | -1.24 | Reduced stability |
| F321C** | <sup>53</sup> | 1 | Probably damaging | -2.5 | Destabilizing | -0.19 | Reduced stability |

These analyses demonstrate the need for an alternative strategy to understand laforin LD patient mutations. Therefore, we designed an experimental pipeline to empirically characterize the range of effects of laforin missense mutations *in vitro* (Fig. 1c). Protein stability, carbohydrate binding, and conformational dynamics were measured using purified proteins, and functional interactions with partner proteins were determined by co-expression in yeast.

### Stability and substrate binding of laforin missense mutations

We previously defined the unique quaternary structure of laforin, a constitutive dimer (Fig. 2a) ^11^. LD missense mutations fall into 4 regions of the laforin crystal structure: the carbohydrate binding module (CBM), the dual specificity phosphatase (DSP) domain, the CBM-DSP interface, and the dimer interface (Fig. 2b). Of the selected mutants, 22/26 could be purified and were utilized for subsequent biochemical analysis. The exceptions were F5S, Y86D, and R108C, affecting core residues of the CBM, and T187A of the DSP, none of which could not be expressed as soluble proteins. The thermal stability of each purified mutant was determined by differential scanning fluorimetry (DSF). We previously demonstrated that short (7 glucosyl units, DP7) and long (24 glucosyl units, DP24) oligosaccharides stabilize laforin, and DSF in the presence of glucans can also be used to assess binding ^11^.

**Fig. 2.**
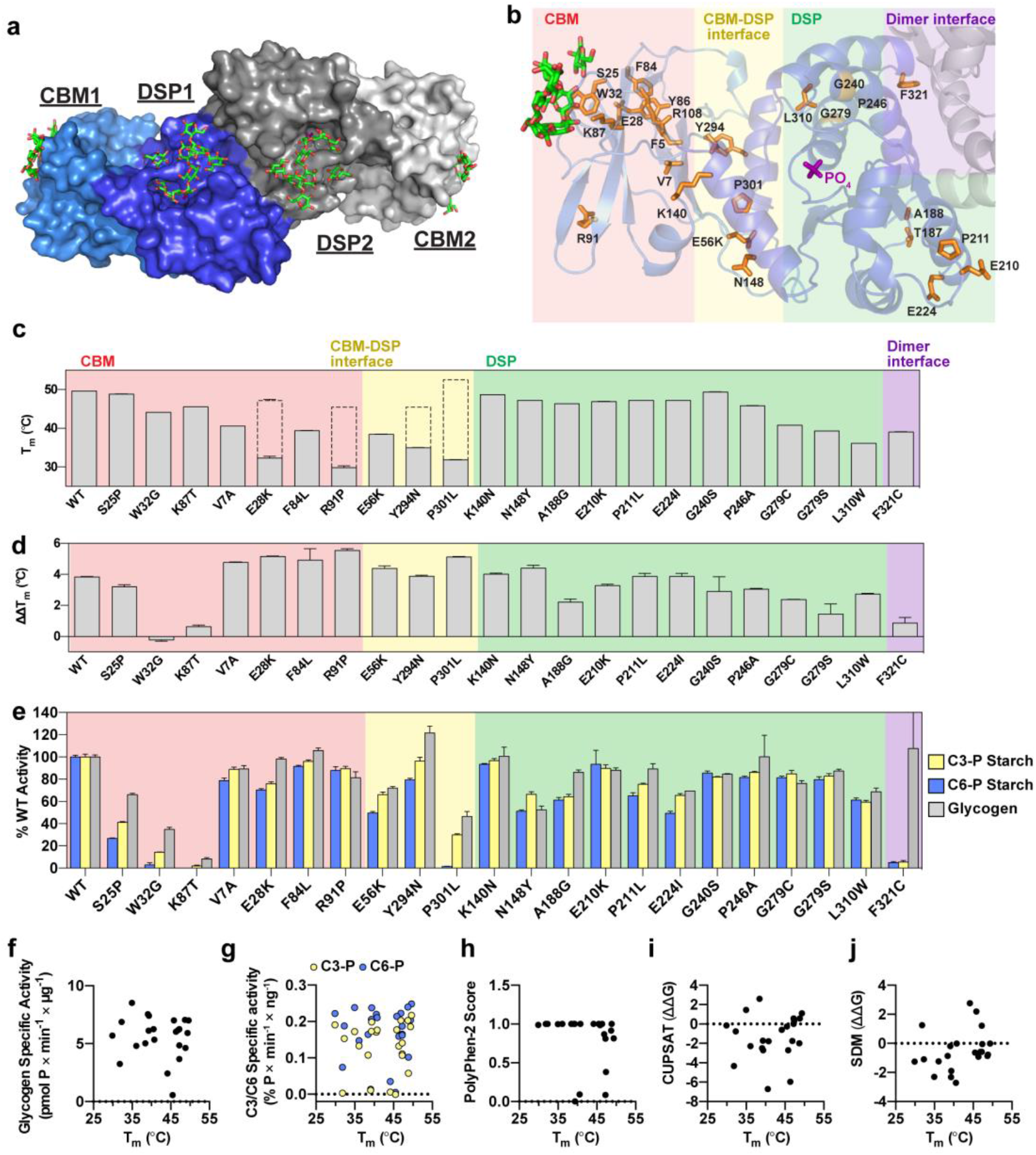
Stability, carbohydrate binding and phosphatase activity of LD mutants. (a) Surface structure of laforin bound to maltohexaose (PDB:4rkk) ^11^. Laforin forms an antiparallel homodimer with a DSP-DSP interface. The CBM and DSP domain of one subunit are shown in light and dark shades of blue, respectively. Bound maltohexaose molecules are shown in green. (b) Ribbon diagram of one subunit of the laforin dimer. Residues affected by missense mutations are shown in orange. Phosphate bound to the active site is shown in purple, and glucans bound to the CBM are shown in green. The CBM and DSP domain are shaded as in (a). Mutations in the CBM are boxed in pink, the CBM-DSP interface in yellow, the DSP in green, and the dimer interface in purple. (c) Mutant stability measured by melting temperature (T_m_). For mutants displaying biphasic melting, the first (primary) peak is represented by the filled grey bar; the dotted bar indicates the approximate T_m_ corresponding to the second peak. (d) The difference in ΔT_m_ displayed for each mutant in the presence of 10 mM DP7 compared to 10 mM DP24, indicated as ΔΔT_m_. (e) Specific activity of laforin mutants with the following substrates: glycogen (grey), C3-P (yellow), C6-P (blue). Activity of each mutant was normalized to WT activity. In (c), (d) and (e), all assays were performed in triplicate, and graphs represent the average ± SD. Also see Figure S2. (f-j) Correlation scatterplots of T_m_ versus glycogen specific activity (i; r=-0.005929, p=0.9786); C3-P and C6-P starch specific activity (j; r=0.2141, p=0.3266 for C3-P; r=0.09289, p=0.6734 for C6-P); PolyPhen-2 score (f; r=-0.4068, p=0.0603), and CUPSAT (g; r=0.2033, p=0.3641) and SDM (h; r=0.3972, p=0.0672) predictions for ΔΔG (kcal/mol).

Wild-type (WT) laforin exhibited a single melting transition and a melting temperature (T_m_) of 49.6°C (Fig. 2c and S1a). DP7 and DP24 stabilized WT laforin (ΔT_m_) by 4.6 and 7.8°C respectively (Fig. S1b). The 3.2°C difference between DP7 and DP24 binding (ΔΔT_m_) reflects the established preference of laforin for long glucan chains (Fig. 2d) ^11,47^. The patient mutations W32G and K87T were slightly destabilized with T_m_ values of 44-45°C (Fig. 2c), but showed minimal shifts of <1°C in the presence of glucans (Fig. 2d and Fig. S1b). These results are consistent with direct glucan engagement by W32 and K87 in the laforin structure ^11^. Mutations in CBM core residues (V7A, E28K, F84L, R91P) and at the CBM-DSP interface (E56K, Y294N, P301L) were highly destabilized with T_m_ values of 40°C or less (Fig. 2c). Notably, in contrast to WT laforin, some of these mutants (E28K, R91P, Y294N, and P301L) displayed a biphasic melting profile with a second T_m_ appearing between 45 and 53°C (Fig. 2c and S1a). Mutations of the CBM core and CBM-DSP interface yielded a greater ΔT_m_ upon addition of glucan, suggesting substrate-induced compensatory stabilization (Fig. S1b). However, these mutants exhibited a preference for long glucans similar to WT laforin (Fig. 2d). The only CBM mutant with no change in stability or binding was S25P (Fig. 2c,d and S1a,b). Mutations of buried DSP residues (G279C, G279S, L310W) were all destabilized with T_m_ values of 36-41°C (Fig. 2c) and showed similar glucan binding as WT (Fig. 2d and S2b). In contrast, DSP mutations (A188G, K140N, N148Y, G240S, E210K, P211L, E224I) showed little effect on either stability or glucan binding (Fig. 2c,d and S2a,b). Finally, F321C affecting the dimer interface showed moderate destabilization and a specific reduction in preferential binding to DP24, as indicated by ΔΔT_m_ values of <1 °C (Fig. 2c,d and S1a,b). We previously demonstrated that while WT laforin is a dimer in solution, F321S and F321C are both monomeric, and dimerization is directly linked to the preferential binding of laforin for long glucans ^11,48^.

### Activity of laforin missense mutations

We next tested the phosphatase activity of the mutants using multiple substrates. First, total phosphate release from laforin’s biological substrate glycogen was determined (Fig. 2e and S1c). Laforin preferentially binds to glucans with long chains, and LBs resemble plant starch in that they are insoluble and contain more phosphate and longer chains than soluble glycogen ^2,37^. Glycogen, LBs and starch are phosphorylated at the C3 and C6 hydroxyls of glucosyl units ^49–51^. Site-specificity assays using radiolabeled starch indicate WT laforin dephosphorylates both positions with a slight preference for C6-phosphate (C6-P) ^52^. Therefore, we tested the phosphatase activity of laforin using C3-P and C6-P labeled starch (Fig. 2e and S1d).

W32G and K87T had significantly impaired phosphatase activity toward all substrates, consistent with their impaired glucan binding (Fig. 2e and S1c,d). Mutations in the CBM-DSP interface had varying effects on glycogen phosphatase activity: Y294N activity was comparable to WT, E56K activity was mildly impaired, and P301L yielded a 50% decrease (Fig. 2e and S1c,d). In the presence of starch substrates, E56K and P301L activities were again impaired, while Y294N had slightly higher activity than WT laforin. Interestingly, C6-P starch dephosphorylation was completely eliminated in P301L, while some C3-P activity was preserved; however, no other mutants displayed a similar imbalance in specificity. Interestingly, the remaining 19 mutants maintained ≥50% of WT activity regardless of substrate, with the exception of F321C (Fig. 2e and S1c,d). F321C displayed unchanged glycogen phosphatase activity but a profound impairment in C3-P and C6-P starch dephosphorylation. The data illustrate a reduced specificity of F321C for LB-like substrates with long glucan chains, consistent with the DSF results (Fig. 2d).

No significant correlation between thermal stability and phosphatase activities was observed (Fig. 2f,g). Furthermore, when we compared the T_m_ of the purified mutants with the *in silico* predictions, no correlation between T_m_ and PolyPhen-2 score or the predicted ΔΔG via CUPSAT or SDM was detected (Fig. 2h,i,j), further emphasizing the need for experimental approaches to define this system.

### Mutation-induced decoupling of the CBM and DSP domains

WT laforin melts with a single sharp peak (Fig. S1a). Most laforin mutants displayed a melting profile of similar shape, even when the curve shifted left due to reduced stability. The exceptional biphasic transitions observed in E28K, R91P, Y294N, and most prominently in P301L suggest decoupling between the CBM and DSP domain in these mutants. To test this hypothesis, we purified the individual laforin domains. The T_m_ value of the CBM, 38.2 °C, and of the DSP domain, 51.7 °C (Fig. 3a), were similar to the T_m_ values of the very prominent P301L transitions, 31.8 and 52.5 °C (Fig. 2c and S1a), supporting our hypothesis. When the CBM and DSP domains were incubated with increasing concentrations of DP7, the CBM exhibited a similar binding curve to full-length laforin, while the DSP domain did not shift even at high concentrations of DP7 (Fig. 3b). These results confirm that the CBM is necessary and sufficient for carbohydrate binding.

**Fig. 3.**
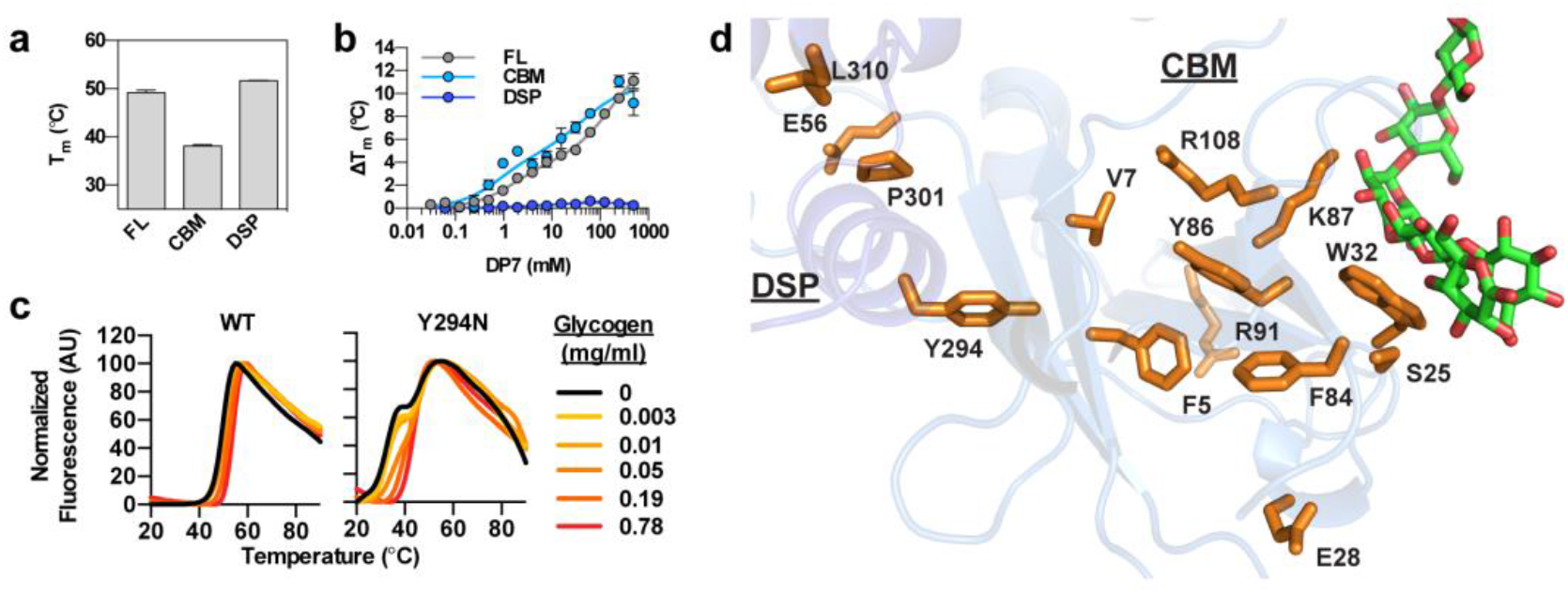
DSF melting profiles of laforin mutants. (a) Thermal stability of CBM and DSP proteins compared to full-length laforin (FL). (b) Binding of FL, CBM and DSP proteins with increasing concentration of DP7. (c) Melting curves of WT and Y294N binding to increasing concentrations of glycogen. (d) The CBM core and CBM-DSP interface is a hotspot for LD mutations. Residues affected by missense mutations are shown in orange. Glucan is shown in green.

The shape of the WT melt curve does not change with glycogen binding (Fig. 3C). However, the melt curve of the decoupled mutant Y294N gradually shifted to a single transition with increasing concentrations of glycogen (Fig. 3c). This phenomenon was also observed with DP7 for the other decoupled mutants (Fig. S1a). These results suggest that glucan binding, even to short glucans, induces a deep integration between the CBM and DSP domain that becomes visible in mutants with CBM-DSP decoupling. Indeed, the CBM core and CBM-DSP interface appears to be a “hotspot” for laforin missense mutations, many of which affect buried residues (Fig. 3d). Furthermore, the mutants with biphasic transitions had an initial T_m_ of less than 37°C (Fig. 2c), indicating they are at a high risk for unfolding at physiological temperatures.

### Solution dynamics of LD mutations determined by hydrogen-deuterium exchange (HDX)

The biochemical effects underlying slow or late-onset cases of LD could not be understood from stability, binding, and phosphatase activity measurements of the associated mutants. In order to further understand the structural and functional perturbations that may alter laforin behavior, we performed hydrogen deuterium exchange (HDX) experiments on a subset of LD mutants. HDX reveals the dynamics of protein conformational states in solution by quantifying exposure to a deuterated solvent over time. We selected key mutants associated with a range of biochemical activities and clinical outcomes: R91P (associated with typical LD; displayed biphasic melting, severe destabilization and no change in activity), G279C (associated with a slower disease progression; destabilized, but no change in glucan binding or activity), P211L (associated with a slower disease progression; no change in stability, binding or activity), and F321C (associated with very late-onset LD; destabilized and has altered specificity for long glucan substrates). 100% peptide coverage for all four mutants was achieved, and peptide profiles between the four mutants were nearly identical (Fig. S2). Residues with a deuteration change of 10% or more were mapped onto the laforin structure (Fig. 4).

**Fig. 4.**
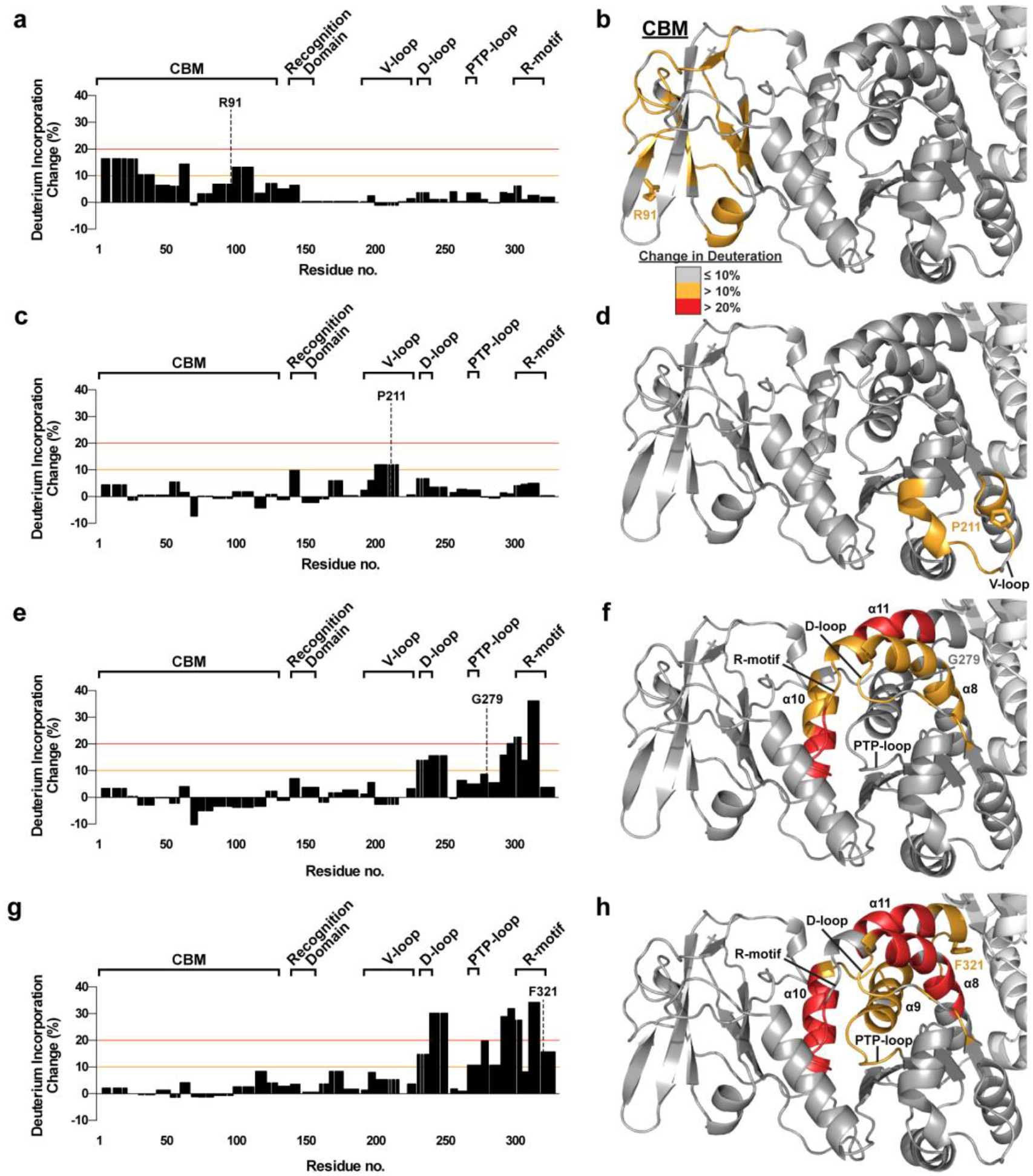
HDX of LD mutants. (a, c, e, g) Change in deuteration incorporation (%) for all peptides are shown. The significance thresholds that were used for mapping onto the laforin crystal structure (b, d, f, h) are marked by orange (10%) and red (20%) gridlines. (b, d, f, h) Deuteration changes induced by mutations are mapped onto one subunit of the laforin structure. The mutated residue is shown in stick representation and labeled. Structural features are labeled in black.

R91P, a mutant which displayed CBM-DSP decoupling by DSF, yielded significant increases in deuteration throughout the CBM (Fig. 4a,b). P211L caused small changes in deuteration with significant increases affecting only the V-loop of the DSP (Fig. 4c,d). G279C also caused significant deuteration increases only in the DSP domain (Fig. 4c,d). Increased solvent accessibility was observed in the D-loop, R-motif, and helices α8 and α11 of the DSP, and α10 at the CBM-DSP interface. F321C affected similar regions of the DSP but to a greater extent (Fig. 4g,h). The catalytic PTP-loop and the adjacent α9 helix also exhibited significantly increased deuteration. The greater solvent accessibility at the DSP-DSP interface is consistent with the loss of dimerization in F321C ^48^.

These HDX data illustrate how LD patient mutations cause local structural perturbations with distinct consequences. R91P induced a general destabilization of the CBM which led to domain decoupling. E28K, which also displayed biphasic melting despite its distance from the CBM-DSP interface, is likely to have conformational changes comparable to R91P. Other mutations in this hotspot that affect the CBM may have similar effects based on their biochemical profiles and location. The HDX results are consistent with an allosteric path between the CBM and DSP domain. These data also align with our previous HDX data revealing conformational changes in the laforin structure induced by glycogen binding.^11^ While P211L induced only local perturbations within the V-loop, G279C and F321C mutations caused greater solvent accessibility throughout other motifs of the DSP domain. HDX data on F321C also support a loss of dimerization in this mutant ^48^. Furthermore, the significant increases in deuteration of α10 in G279C and F321C further support the importance of CBM and DSP integration for laforin function.

### Effects of LD mutations on protein-protein interactions

We have shown that some LD mutants are pathogenic because they impair glucan binding (W32G and K87T) or have extremely low thermal stability (E28K, R91P, Y294N, and P301L). F321C only impairs preferential binding to long glucans and is associated with very late onset ^53^. However, many LD mutants maintain phosphatase activity and display little to no destabilization, indicating there are additional disease-relevant functions of laforin. Laforin is known to interact with multiple proteins involved in glycogen metabolism. Therefore, to understand the effects of patient mutations on laforin interactions in a cellular context, we assessed the impact of laforin mutants with other proteins known to interact with laforin.

Malin and Protein Targeting to Glycogen (PTG) are well-characterized laforin binding partners ^12,13,54-57^. To measure their interaction with laforin, we employed a directed yeast two-hybrid assay that has previously been used to identify and define laforin binding partners ^12,13,57-59^. WT laforin and mutants were expressed as bait proteins fused with the DNA-binding domain of the LexA protein. WT malin and PTG were expressed as prey fused with the Gal4 activation domain (GAD) and β-galactosidase (i.e. β-gal) activity was used to quantify interaction between bait and prey.

Strikingly, a significant number of laforin mutations reduced or abolished interaction with either PTG, malin, or both (Fig. 5a). This finding highlights the importance of laforin as a glycogen-associated protein scaffold. Interestingly, mutations at the CBM glucan binding site, W32G and K87T, reduced PTG interaction, while not affecting the interaction with malin (Fig. 5a). In contrast, F321C specifically abolished the malin interaction (Fig. 5a). S25P, which had no obvious biochemical defect, lost interaction with both malin and PTG (Fig. 5a). Additionally, we noticed a spatial pattern to these interaction data: mutations at the CBM specifically affected the PTG interaction, mutations at the dimer interface specifically affected the malin interaction, and centrally located mutations affected both interactions. Therefore, we hypothesized that the binding sites for PTG and malin are spatially distinct. This arrangement is logical since PTG has been shown to be a substrate for malin in a laforin-dependent fashion, and therefore both PTG and malin likely interact with laforin at the same time ^54^. We tested this hypothesis by performing yeast triple hybrid experiments in which laforin was expressed in the presence of both malin and PTG. One partner was expressed as prey, fused to GAD, while the other was expressed with only an HA tag. If the two binding partners had overlapping binding sites, the HA-tagged partner would displace the first and result in reduced β-gal activity. No change in β-gal activity was observed when HA-tagged PTG was expressed in addition to LexA-laforin and GAD-malin (Fig. 5b). To account for the possibility that the laforin-malin interaction was too strong for malin to be displaced by PTG, we also performed the reverse experiment in which HA-tagged malin was expressed in addition to LexA-laforin and GAD-PTG. Again, no change was observed in β-gal activity (Fig. 5b). These data strongly support the hypothesis that PTG and malin have independent binding sites on laforin, and these binding sites are differentially affected by LD missense mutations due to local structural perturbations.

**Fig. 5.**
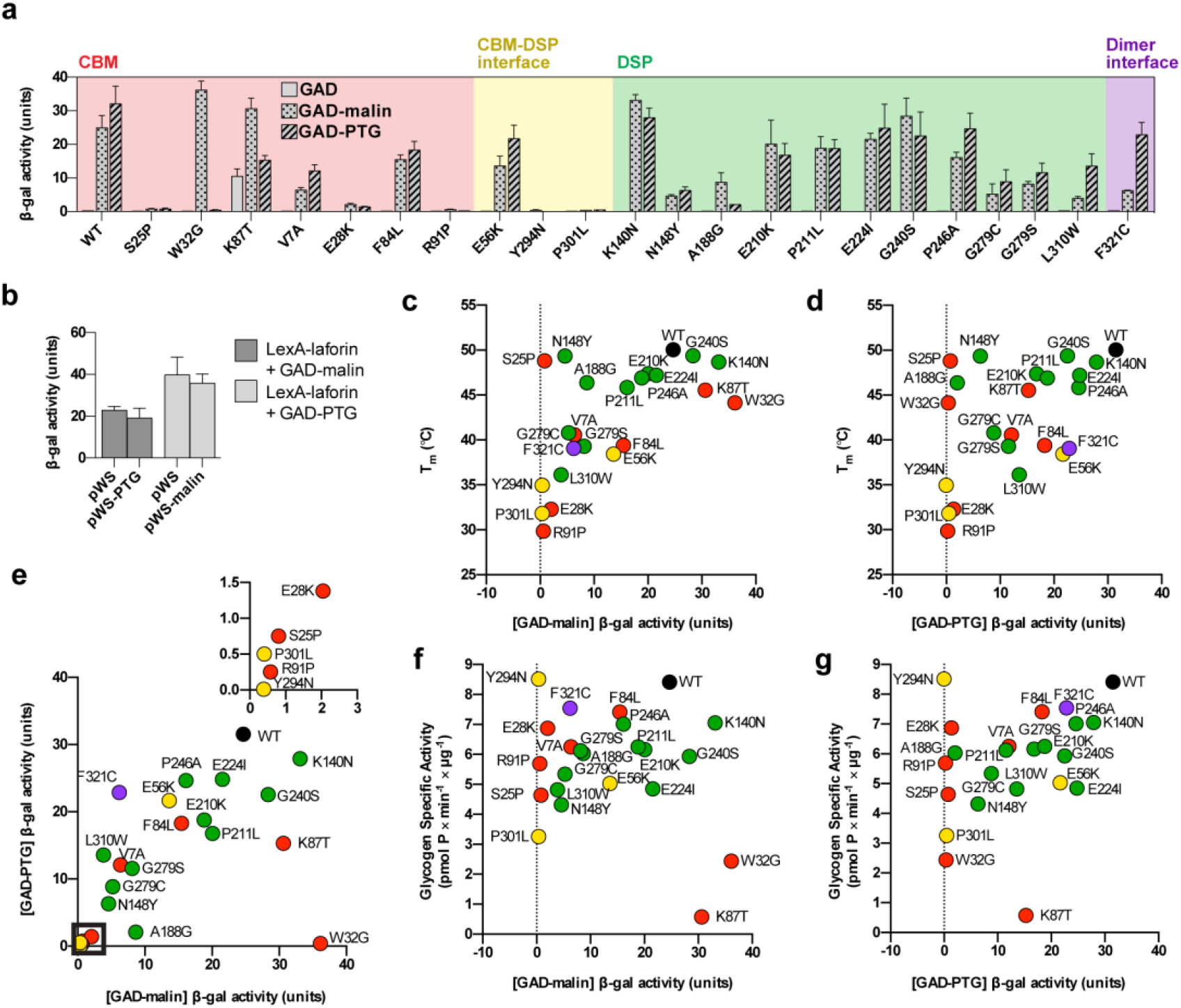
Effects of LD mutants on the interaction of laforin with malin and PTG. (a) Interaction of WT laforin and mutants with GAD, GAD-PTG, and GAD-malin fusion proteins. Mutants are color coded by class as in Figure 2. (b) Triple hybrid experiment demonstrating noncompetitive binding of malin and PTG to laforin. LexA-laforin was expressed with GAD-malin and pWS-PTG or GAD-PTG and pWS-malin. (c, d, e, f, g) Correlation between T_m_ and GAD-malin interaction (c; r=0.6087, p=0.0021), T_m_ and GAD-PTG interaction (d; r=0.499, p=0.0154), GAD-malin and GAD-PTG interactions (e; r=0.6462, p=0.0009), glycogen activity and GAD-malin interaction (f; r=0.01087, p=0.9607) and glycogen activity and GAD-PTG interaction (g; r=0.3646, p=0.0872). In (e), box corresponds to inset. Color of dots correspond to mutation location as in Figure 2: CBM (red), CBM-DSP interface (yellow), DSP (green), dimer interface (purple).

### Correlation of defects in Laforin

The effects of LD mutations on multiple aspects of laforin have now been defined to alter stability, glucan binding, substrate specificity, dephosphorylation, and interaction with established binding partners. To determine whether these aspects of laforin function were correlated, we performed pairwise correlation analyses between T_m_, ΔΔT_m_, specific activities, PTG interaction, and malin interaction measurements (Fig. S3 and Table S1). A significant positive correlation was observed between the malin interaction and T_m_ (Fig. 5c, p=0.0021). A positive correlation between PTG interaction and T_m_ was also observed (Fig. 5d, p=0.0154). A very strong correlation was identified between PTG and malin interaction (Fig. 5e, p=0.0009). No significant correlation was found between glycogen specific activity and malin or PTG interaction (Fig. 5f,g). As expected, strong correlations were also found between glycogen and starch activities, and between C6-P starch activity and ΔΔT_m_, since starch dephosphorylation requires substrate specificity for which ΔΔT_m_ is a proxy (Fig. S3 and Table S1). In conclusion, these data illustrate that LD mutants primarily affect laforin’s ability to bind to glucan substrates or protein binding partners; and protein binding is dependent on structural integrity of laforin within the interacting region. Additionally, mutants fall into particular patterns of functional effects related to their location within the structure.

## DISCUSSION

In the present study, we presented four LD clinical cases with different outcomes and defined the molecular defects of *EPM2A* missense mutations affecting laforin function. Genotypephenotype correlations in the LD population are extremely difficult due to a small number of patients, high allelic heterogeneity, frequent under-diagnosis, and limited clinical data. We employed an integrated experimental pipeline and define mutation pathogenicity for 26 patient mutations using biochemical methods.

### Functional classes of *EPM2A* missense mutations

We propose a classification of laforin mutants based on their biochemical profile (Table 3). Class I mutations directly affect CBM carbohydrate binding (Table 3). The CBM is essential for carbohydrate binding and W32 and K87 are primary residues involved in glucan interaction. Class I mutations are only slightly destabilizing to the laforin structure, and their phosphatase activity is greatly reduced due to reduced substrate affinity. These mutations also abolish the laforin interaction with PTG. Although both W32G and K87T still interact with malin, they cannot bind glycogen and thus cannot target malin to glycogen-associated substrates. Based on our biochemical results, it is reasonable to conclude that since these mutations are highly detrimental to all aspects of laforin function they lead to a severe phenotype with rapid progression, as we observed in patients 2 and 3 (Table 1).

**Table 3.**
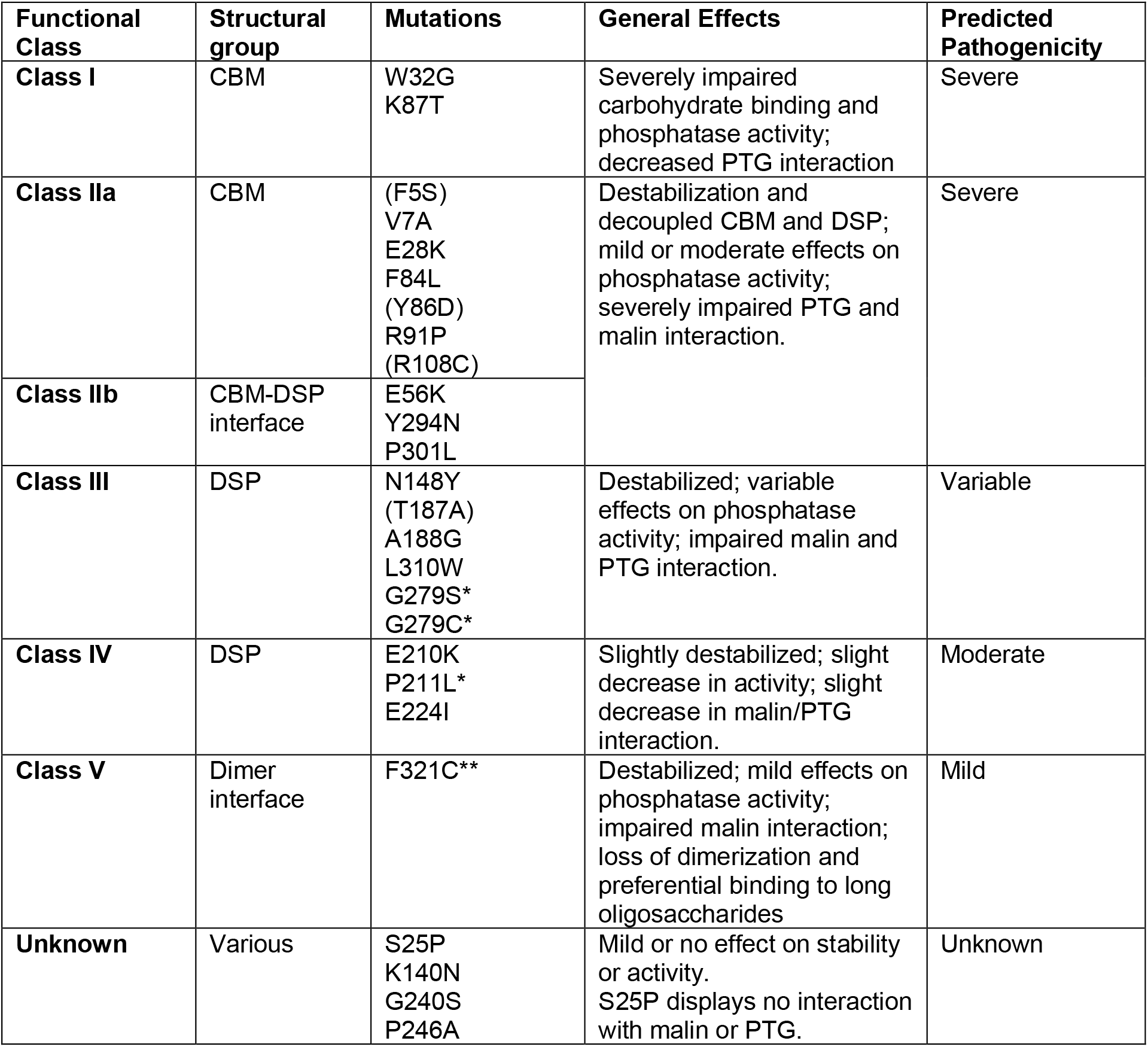
Classification of LD-causing *EPM2A* missense mutations based on biochemical and clinical data. Asterisk(s) indicates this mutation has been associated with a milder phenotype, either in the compound heterozygous (*) or homozygous (**) state. Mutants in parentheses are likely classifications but were not fully characterized due to extensive proteolysis.

Class IIa mutations affect the CBM core and Class IIb mutations affect the CBM-DSP interface (Table 3). Although structurally distinct, Class IIa and IIb mutations produce similar biochemical defects: they dramatically destabilize laforin and impair interactions with both PTG and malin. Most of these mutations have little or no effect on phosphatase activity, with the exception of P301L, which may have compromised catalytic activity due to its severe instability. Patients either homozygous or compound heterozygous for Class II mutations have been reported to exhibit a classic clinical course, such as patient 1 in the present report, who was compound heterozygous for the E28K mutation. Class II mutations may also be associated with early-onset LD because of their severe effects on laforin stability and binding to other proteins. A family of three children all homozygous for R108C experienced seizure onset at ages 5, 8 and 10 with early onset learning disability that manifested at 4 years old ^60^. Another patient homozygous for Y86D displayed early onset learning problems ^40^. Y86D and R108C were two of the four mutations that could not be expressed and purified, likely due to very severe stabilization. Based on their location within the CBM, these mutants are likely to have similar biochemical effects and clinical outcomes as other Class IIb mutants.

Class III mutations affect the DSP domain (Table 3) and have variable effects on laforin stability, PTG and malin interactions. All Class III mutations have mild or moderate effects on phosphatase activity, and they do not affect carbohydrate binding. For LD patients, Class III mutations are associated with variable disease progression ^46,61^. These mutations also significantly impair the interaction with PTG and malin. We report that G279C is associated with a slower clinical course in a patient, which is consistent with previous reports of patients carrying the G279C or G279S mutation. One patient (G279C/R241X) did not present with seizures until the age of 21 years old, and cognitive impairment did not develop until the patient was 24 years old ^40^. Another patient (G279S/R241X) had seizure onset at age 17 but lived beyond the age of 40 ^32^. Since Class III mutations retain some interaction with malin and PTG, they may delay LD onset. We recently published a study on a compound heterozygous patient carrying the missense mutation N163D and the nonsense mutation Y112X ^62^. This patient had her first seizure at 16, and at 28 years old she was still very cognitively engaged and able to walk. We showed that N163D had no effect on laforin stability, carbohydrate binding, or activity, but impaired the interaction of laforin with its binding partners. N163 is in the DSP domain and the biochemical profile of the N163D mutation is consistent with other mutations in Class III (Table 3). Therefore, mutations in class III can produce both atypically moderate and typically severe phenotypes.

Class IV mutations affect a surface-exposed region of the V-loop adjacent to the dimer interface (Table 3). These mutations display no major defects in laforin stability, carbohydrate binding, phosphatase activity, or interaction with PTG or malin. Laforin interacts with additional proteins, including AMP-activated protein kinase (AMPK), glycogen synthase, and R6, another regulatory subunit of PP1 ^63–66^. This region is likely important for binding to one or more of these additional proteins. Therefore, since class IV mutations yield a more moderate effect on laforin function, they may be generally associated with a less severe disease progression.

Class V mutations affect laforin dimerization and therefore substrate specificity. F321 mutations render laforin monomeric and decrease specific binding to substrates with long glucans like LBs. The reduced stability of these mutations is likely a result of dimerization loss. F321 mutations do not affect the interaction of laforin with PTG, but they reduce the malin interaction. F321C is associated with a homozygous case of extremely mild LD in which the patient had a history of seizures that were relatively controlled with sodium valproate and lived to the age of 56 ^53^. The biochemical effects of F321C and the extremely unusual clinical outcome are so unique that a separate class is merited, but the class could easily expand to include other mutants with similar effects.

While these classes explain the defect induced by the majority of patient mutations analyzed, no clear defect was identified in K140N, G240S, P246A, and the reason for the impaired interaction of S25P with malin and PTG is unclear. Thus, there are additional unidentified factors causing pathogenicity, and these mutants have not yet been classified. It is possible that some pathogenic mutations alter post-translational modifications of laforin. An *in vitro* study showed that laforin is phosphorylated at S25 by AMPK, and this phosphorylation enhances the malin-laforin interaction in yeast. This result could explain the absence of both malin and PTG interaction in our study, but the hypothesis requires further validation. There is strong *in vitro* evidence that laforin is ubiquitinated by malin ^12,13,56^. While the specific lysine residue(s) that are ubiquitinated have not been identified, only 3 of the 11 laforin lysine residues are surface exposed: K140, K219, and K323. It is possible that K140 is a primary ubiquitination site and that impaired laforin ubiquitination leads to LD by a yet unknown mechanism. G240S and P246A are centrally located in the DSP between the active site and dimer interface. These mutations have little to no effect on stability, glucan binding or interaction with PTG or malin; therefore, like Class IV mutations, they may affect interactions with other laforin substrates.

### Classifying other known and novel *EPM2A* missense mutations

Based on our study, known and novel missense mutations require biochemical profiling for stability, glucan binding, glucan specificity, and protein-protein interactions. Since phosphatase activity is not correlated with any of these functions and is preserved in most pathogenic mutants, it is not a key readout in our classification. We propose that mutation classification requires five or fewer steps (Fig. 6a). (1) If a mutant strongly impairs or abolishes glucan binding, it would be a Class I mutant and therefore severely pathogenic. Since the CBM is necessary and sufficient for glucan binding, these mutations would typically localize to the CBM. (2) If a mutant specifically impairs binding to long glucans, then it likely also impairs dimerization, and vice versa. These mutants would be in Class V, and likely more mildly pathogenic. (3) Mutants that are significantly destabilized and exhibit impaired interactions with both malin and PTG would fall into Class II or III. (4) Further division into Class IIa, IIb or III would be based on the location of the mutation within the laforin structure. Class II mutations are more commonly severe, with Class III ranging in severity. (5) Remaining mutants lacking defects in glucan binding, stability, or malin/PTG interaction would only be classified into Class IV if they affect the V-loop, which would likely impact a protein-protein interaction site. Mutations in Class IV may be milder with respect to patient progression. If the mutation is not located in the V-loop, further analysis would be required to understand if it affects a post-translational modification, impairs interaction with another binding partner, or causes some other defect. As these analyses are further developed, additional classes may be added to capture additional aspects of laforin function and mutant pathogenicity.

**Fig. 6.**
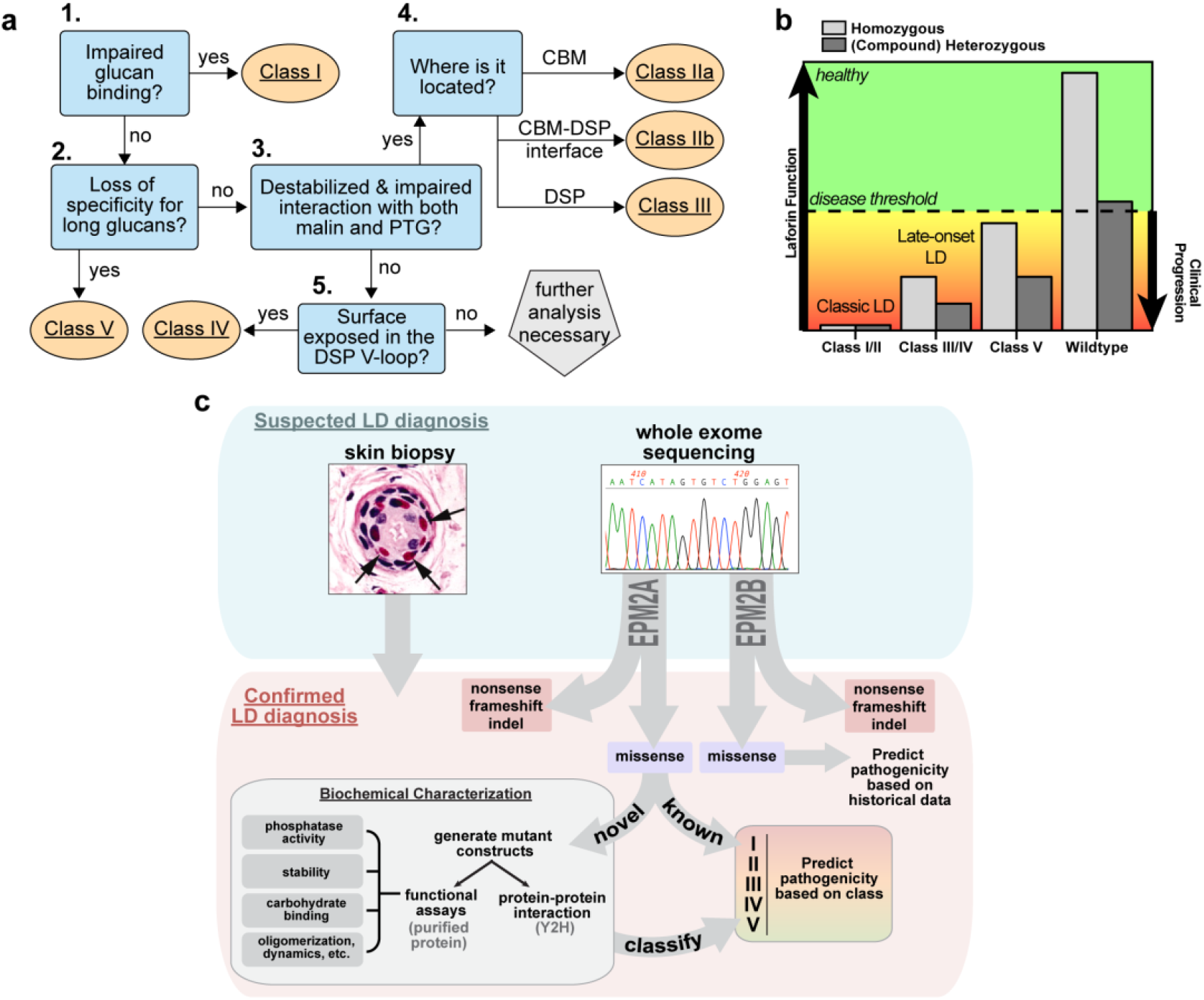
An empirical pipeline to facilitate LD personalized medicine. (a) Classification of a missense mutation requires only five steps. (b) LD progression is dependent on the type of mutation(s) carried by LD patients. In individuals who are homozygous or heterozygous for the wild-type allele, i.e. carriers and non-carriers, laforin function is above the disease threshold and therefore they are healthy. LD patients with Class I or Class II mutations will likely experience classic LD progression. This is likely to be the case whether the patients are homozygous for these missense mutations or complex heterozygotes for missense and deleterious mutations (such as nonsense or indels). LD patients with Class III or Class IV mutations will likely experience a slower progression; particularly if they are homozygous for these mutations. LD patients with Class V mutations will exhibit the slowest progression. (c) A suspected LD diagnosis will be confirmed by genetic testing. Skin biopsies should also be performed to facilitate comparisons of LB load between patients. Nonsense, frameshift, or indel mutations in either *EPM2A* or *EPM2B* would be complete loss-of-function mutations and the most pathogenic. If known loss-of-function or *EPM2A* missense mutation(s) are identified, patient progression could be predicted immediately. If novel *EPM2A* missense mutations are identified, mutants could be characterized and classified within a matter of weeks in order to predict clinical progression for that patient.

### Late-onset LD and genetic effects

An obvious additional complicating factor for predicting clinical progression is that many patients are compound heterozygotes (carrying two different chromosomal aberrations that affect the same gene). Since LD is a recessive disease, it is likely that a disease threshold exists regarding decreased laforin function (Fig. 6b). Our data indicate that laforin function most relevant to LD progression is its ability to coordinately bind glycogen, malin and other glycogen-associated proteins. When laforin function is above a specific threshold then individuals are completely healthy, e.g. heterozygous LD parents. Below this threshold, there is likely a gradation of how rapidly patients progress down a clinical LD path. Both the type of mutation and whether the mutation is in the homozygous or heterozygous state influences disease progression. For example, the slowest onset form of LD associated with an *EPM2A* mutation was reported in a patient homozygous for the Class V mutation F321C ^53^. In contrast, compound heterozygous patients carrying a class III or IV mutation in addition to a deleterious mutation such as a nonsense mutation or indel would experience a clinical course with slower progression than classic LD and yet still more rapid than patients with class V mutations ^40,62^. The most severe and rapidly progressing LD cases are homozygous or compound heterozygous patients with class I or II mutations leading to nonfunctional protein. Patients with these mutations are likely indistinguishable from patients with only nonsense mutations and/or indels.

In addition to the class of mutation, modifier effects are certain to play a role in clinical manifestation. These effects could be either genetic or environmental. Patients carrying the same mutations in ethnic isolates show some phenotypic variability ^67^, and siblings carrying identical mutations sometimes display differences in disease progression, suggesting a role for genetic factors ^40,60,68,69^. For example, a PTG variant has been reported to contribute to a slower disease course ^70^. Differences in medical care can also influence clinical progression ^30^. However, a clinically homogeneous patient progression is often reported within families and among genetic isolates ^71,72^.

### An empirical pipeline for personalized medicine

We are currently expanding our analysis to include all known *EPM2A* missense mutations. In the future, when genetic testing confirms an LD diagnosis and reveals a novel *EPM2A* missense mutation, it could be quickly biochemically assayed and classified (Fig. 6c). If a known *EPM2A* missense mutation is detected, no assays would be required since the biochemical profiles will already be known. Skin biopsies should also be performed to supplement the diagnosis and determine whether LBs enrichment correlates with disease severity or progression. Although biochemical studies have not yet been performed for *EPM2B* missense mutations, case studies indicate specific *EPM2B* mutations can also cause late-onset or slow LD ^32,73^. This empirical pipeline would permit clinicians to make a more accurate prognosis for LD patients based on the mutations they carry.

Genetic screening during pregnancy or at birth is already widely employed to identify genetic diseases and chromosomal abnormalities ^74^. Life-threatening disorders, even those as rare as LD, may soon be added to these early genetic screens and more cases of mild or late-onset LD will be identified. Pre-symptomatic detection of LD would be extremely valuable since early treatment will better ameliorate and could even prevent LD. However, benign polymorphisms also need to be correctly differentiated from mild, moderate and severely pathogenic mutations to prevent a false diagnosis and/or unnecessarily aggressive treatment. Our data reveal distinct characteristics of pathogenic mutations, differentiate mild from severe mutations, and may facilitate differentiation of benign polymorphisms from pathogenic mutations to prevent false diagnosis.

Excitingly, preclinical studies of LD therapeutics are currently underway with multiple lines of treatment in development ^34–36^. The first published therapeutic strategy utilizes an antibody-enzyme fusion that degrades LBs, the toxic carbohydrate aggregates that cause LD ^37–39^. This drug is administered directly into the CNS, eliminating brain LBs and correcting the cerebral metabolic phenotype in LD mouse models. Another promising treatment in development involves antisense oligonucleotides to downregulate glycogen synthase expression, which halted LB formation in a preliminary murine study ^35^. A third strategy utilizes small molecules to inhibit glycogen synthase activity ^75^. A final strategy consists of repurposing approved drugs such as metformin with improved outcomes in mouse models ^76,77^. With the large number of novel mutations arising in new LD patients, our biochemical pipeline will be extremely useful for providing patients with a personalized diagnosis and treatment strategy once therapies become clinically available.

## EXPERIMENTAL PROCEDURES

### Cloning, protein expression, and protein purification

All pET28b and pEG202 laforin mutants were generated by site-directed mutagenesis (QuickChange Lightning, Agilent; Q5 Site-Directed Mutagenesis, New England BioLabs; GENEWIZ Site-Directed Mutagenesis). All pET28b mutants were expressed in BL21-Codon Plus *E. coli* cells and purified using immobilized metal affinity chromatography and a Profinia Purification System (BioRad) and size exclusion chromatography via an ÄKTA fast protein liquid chromatography system (GE Healthcare). Purity of proteins was determined by SDS-polyacrylamide gel electrophoresis (PAGE) with Coomassie staining.

### Differential scanning fluorimetry (DSF)

Experiments were performed using a CFX96 Real-Time PCR system (BioRad). Individual reactions contained 2 μM protein and 5X SYPRO Orange Protein Gel Stain (Invitrogen). DP7/maltoheptaose (Elicityl), DP24 maltodextrins (Elicityl), or rabbit liver glycogen (Sigma) were used as substrates in DSF reactions. Melting was monitored from 20 to 90°C at a ramp rate of 1°C/50sec. Melting temperature (T_m_) was calculated from a Gaussian fit of the first derivative of the melting curve. Data analyses and binding fits were determined using the Prism software (Graphpad).

### Glycogen dephosphorylation assays

Glycogen was purified from rabbit muscle as previously described ^10,50^. Phosphate release was quantified using the Pi ColorLock Gold Phosphate Detection system (Innova Biosciences), a commercial reagent based on the malachite green assay for detecting inorganic phosphate ^10,78^. Kinetics of dephosphorylation were confirmed as previously reported ^10^. Assays were performed in 100 μL reactions containing 2.5 μg enzyme, 2 mM DTT, and phosphatase buffer (100 mM sodium acetate, 50 mM bis-Tris, 50 mM Tris-HCl, pH 6.5) at 25°C. Mutants were assayed in the linear range with respect to time, enzyme amount, and substrate concentration. For specific activity determinations, reactions were performed for 30 minutes with 10mg/ml glycogen.

### Site-specific dephosphorylation assays

Radiolabeled starch was prepared as previously described ^79–81^. Briefly, phosphate-free *Arabidopsis (sex1-3)* starch was phosphorylated with purified glucan water dikinase (GWD) and phospho-glucan water dikinase (PWD). The C6-labeled starch was prepared by including [β ^33^P]-ATP during the GWD incubation and unlabeled ATP during the PWD incubation. The C3-labeld starch was prepared by including unlabeled ATP during the GWD incubation and [β ^33^P]-ATP during the PWD incubation. Labeled starch was washed thoroughly after each phosphorylation step to remove unbound phosphate. [β ^33^P]-ATP was obtained from Hartman Analytic.

Dephosphorylation reactions were performed in a volume of 150 μL with 50 ng of enzyme in dephosphorylation buffer (100 mM sodium acetate, 50 mM bis-Tris, 50 mM Tris-HCl, pH 6.5, 0.05% [v/v] Triton X-100, 1 μg/μL [w/v] BSA, and 2 mM DTT) and 3 mg/ml of either C6- or C3-labeled starch. After 2.5 minutes on a rotating wheel at 25°C, reactions were quenched by adding 50 μL of 10% SDS, and then centrifuged for 5 minutes at 13,000 rpm to pellet the starch. 150 μL of the supernatant was added to 3 mL scintillation liquid, and ^33^P release was quantified using a 1900 TR liquid scintillation counter (Packard).

### Yeast two-hybrid assays

For yeast two-hybrid assays, pEG202-laforin encoding a LexA-laforin fusion protein and pACT2, pACT2-malin and pACT2-PTG encoding Gal4 activation domain (GAD) and GAD fusions have been previously described ^57,59,62,70^. pWS-malin and pWS-PTG encoding HA-tagged proteins were used for the triple hybrid assay. *Saccharomyces cerevisiae* THY-AP4 (*MATa, ura3, leu2, lexA::lacZ::trp1, lexA::HIS3, lexA::ADE2*) were transformed with the indicated plasmids, and transformants were grown in selective SC medium. Yeast two-hybrid assays were performed as previously described ^62^. Briefly, transformants were screened for β-galactosidase activity using a filter lift assay. The strength of the interaction was determined by measuring β-galactosidase activity in permeabilized yeast cells and expressed in Miller units.

### Hydrogen deuterium exchange (HDX)

Quenching experiments for laforin have been previously described ^11^. Deuterium exchange was performed at various time points (10-100,000sec), followed by pepsin digestion and analysis on an Orbitrap Elite mass spectrometer (Thermo Fisher Sci) (See Supplemental Methods). Proteome Discoverer software (v1.3, Thermo Scientific) was used to identify the sequence of the digested peptide ions from their MS/MS data. HDXaminer (Sierra Analytics, Modesto, CA) was utilized to confirm the peptide identification and calculate the centroids of isotopic envelopes of all the peptides. The level of deuterium incorporation of each peptide was calculated by applying back-exchange correction ^82^. The ribbon maps (Supplemental File 1) were generated from deuteration level of overlapping peptides to improve the resolution of the HDX data, and difference maps (Supplemental File 2) show changes in mutants compared to WT laforin.

### Structural and statistical analysis

PyMol 2.0 was used for structural analysis and generating molecular graphics ^83^. All statistical tests were performed using Prism Software (GraphPad). Correlation coefficients were determined using the nonparametric Spearman correlation.

## AUTHOR CONTRIBUTIONS

M.S.G. and C.W.V.K. conceived the study. M.M.C. and J.S. collected and analyzed clinical data. M.K.B. and J.W. generated mutants. M.K.B., M.S.G., C.W.V.K., Z.S., S.S., and J.W. purified proteins, performed assays, and analyzed data. R.V., M.A.G.G., and P.S. performed Y2H experiments. S.L. performed HDX experiments and analyzed data. M.K.B., C.W.V.K. and M.S.G. wrote the paper.

## ACKNOWLEDGEMENTS

This work was supported by the National Institutes of Health [R01 NS070899 to M.S.G., P01 NS097197 to M.S.G., R35 NS116824 to M.S.G., and F31 NS093892 to M.K.B], the National Science Foundation [DBI2018007 to M.S.G.], and an Epilepsy Foundation New Therapy Commercialization Grant to M.S.G. M.K.B. has received funding from the European Union’s Horizon 2020 research and innovation programme under the Marie Skłodowska-Curie grant agreement [No. 754510M]. This project also received funding from the Spanish Ministry of Economy and Competitiveness [SAF2017-83151-R to P.S. and RTI2018-095784b-100SAF to J.M.S].

