## Supplementary Material for "An empirical pipeline for personalized diagnosis of Lafora disease mutations"

### SUPPLEMENTAL INFORMATION

Supplemental file 1. Ribbon maps of HDX results for WT laforin and LD mutants.
Supplemental file 2. Difference maps of HDX results for LD mutants compared to WT laforin.

**Supplemental Methods**

Prior to carrying out hydrogen-deuterium exchange (HDX) experiments, the quenching condition for optimal sequence coverage of laforin was established as previously described[1] as 0.08M GuHCl, 0.1M Glycine, 16.6% Glycerol, pH 2.4. Functional HDX experiments were initiated by dilution of 3 μl of stock solution (laforin WT or laforin mutants at 1 mg/ml) into 9 μl of $D_2O$ buffer (8.3 mM Tris, 150 mM NaCl, $pD_{READ}$ 7.2) and incubate at 0ºC. The exchange reactions were quenched after various exchange time-points (10, 100, 1000, 10,000 and 100,000 sec) by the addition 18 μl of the optimal quench solution and the quenched samples were flash frozen with dry ice. Un-deuterated and equilibrium-deuterated control samples were also prepared as previously described [2]. All frozen samples were pass over an immobilized pepsin column (16 μl) at a flow rate of 25 μl/min and digested peptides were collected on a C18 trap column (Optimize Tech, Opti-Trap, 0.2x2 mm) for desalting. The peptide separation was performed on a C18 reverse phase column (Agilent, Poroshell 120, 0.3x35 mm, 2.7 μl) with a linear gradient of 8-48% B over 30 min (A: 0.05% TFA in $H_2O$; B: 80% acetonitrile, 0.01% TFA, and 20 % $H_2O$). MS analysis was performed on the Orbitrap Elite mass spectrometer (Thermo Fisher Sci), which was adjusted for HDX experiments [3]. The resolution of the instrument was set at 120,000 at m/z 400.

### Supplemental Figures

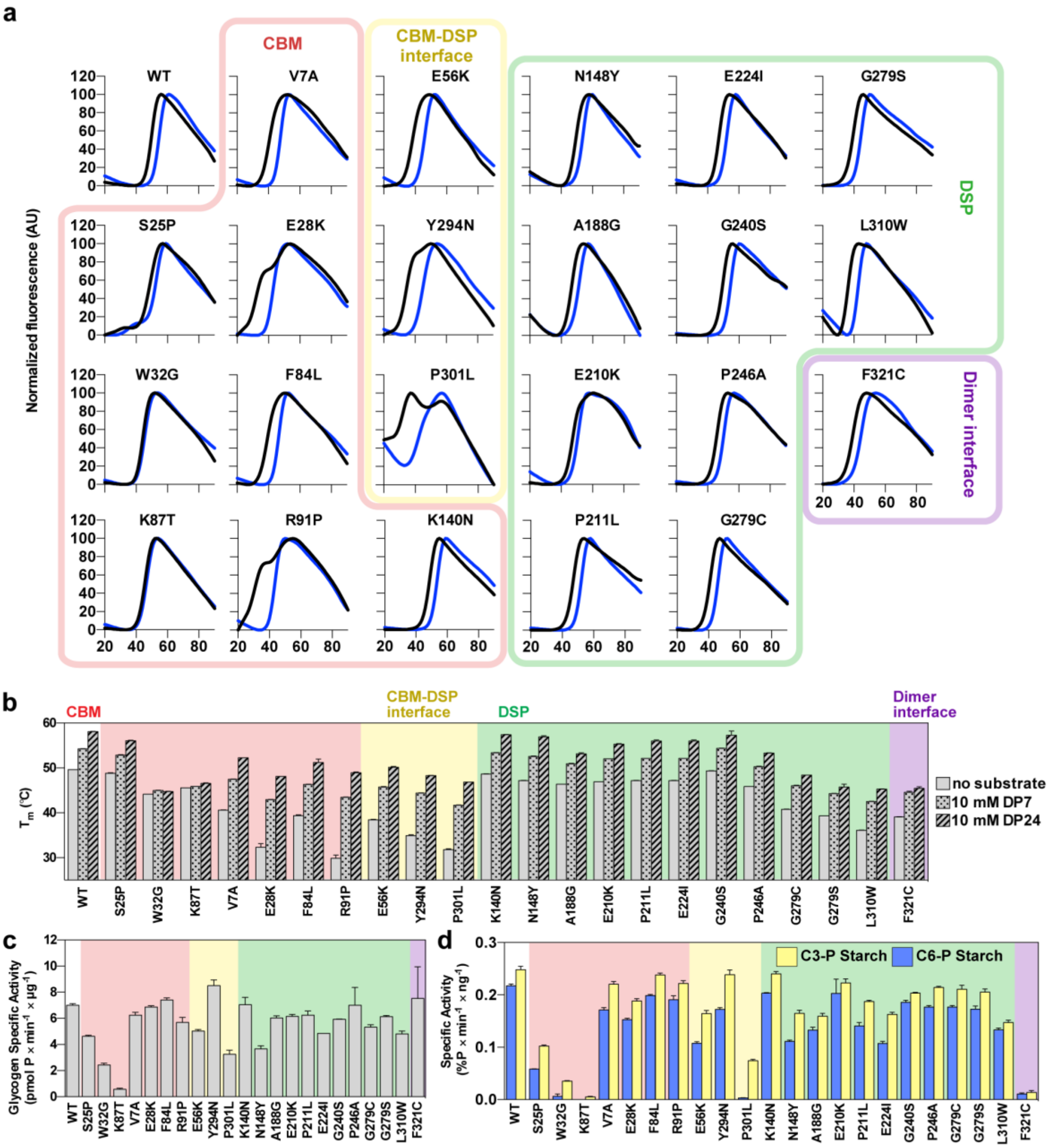


**Fig. S1. Stability, carbohydrate binding, and specific phosphatase activity.** (a) Melting profiles of proteins in the absence of substrate are shown in black. Melting profiles in the presence of 10 mM DP7 are shown in blue. Data are representative of 3 replicates. (b) Absolute $T_m$ for mutants without substrate and in the presence of 10 mM DP7 or 10 mM DP24. For mutants with biphasic melting, only the $T_m$ corresponding to the first transition is shown. (c) Specific glycogen activity of LD mutants. (d) Specific activity of LD mutants with C3-P and C6-P starch substrates.

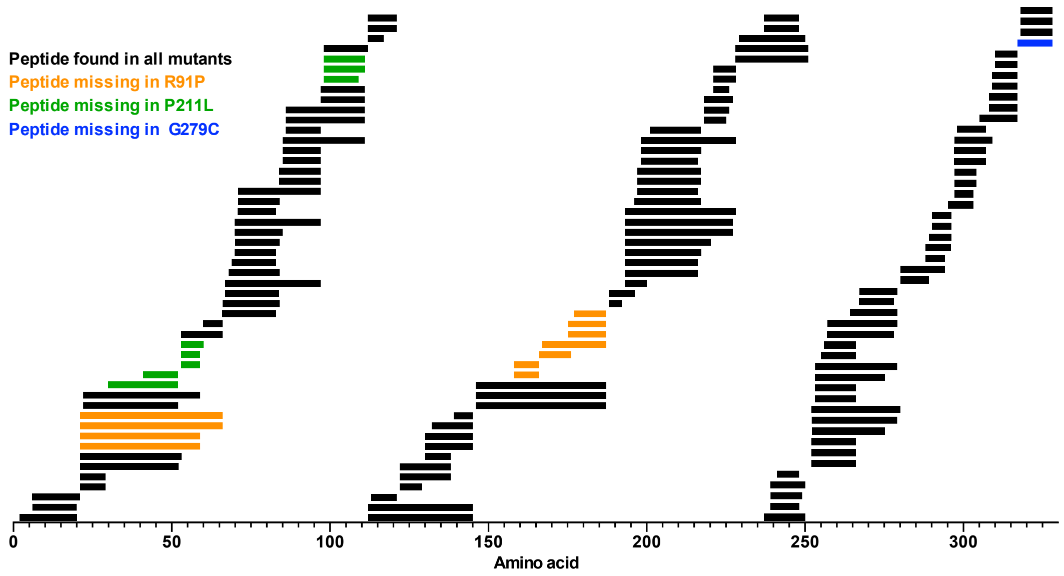


**Fig. S2. HDX peptide coverage of laforin mutants.** Sequence coverage of laforin by pepsin digested peptides. 100% of residues 2-228 were covered by at least one peptide. A total of 148 high-quality peptides were identified. Not all mutants contained every peptide: R91P lacked 11 (orange), P211L lacked 8 (green), and G279C lacked 1 (blue). F321C contained all peptides. The slight changes in pepsin digestion patterns were due to the structural perturbations induced by the mutations.

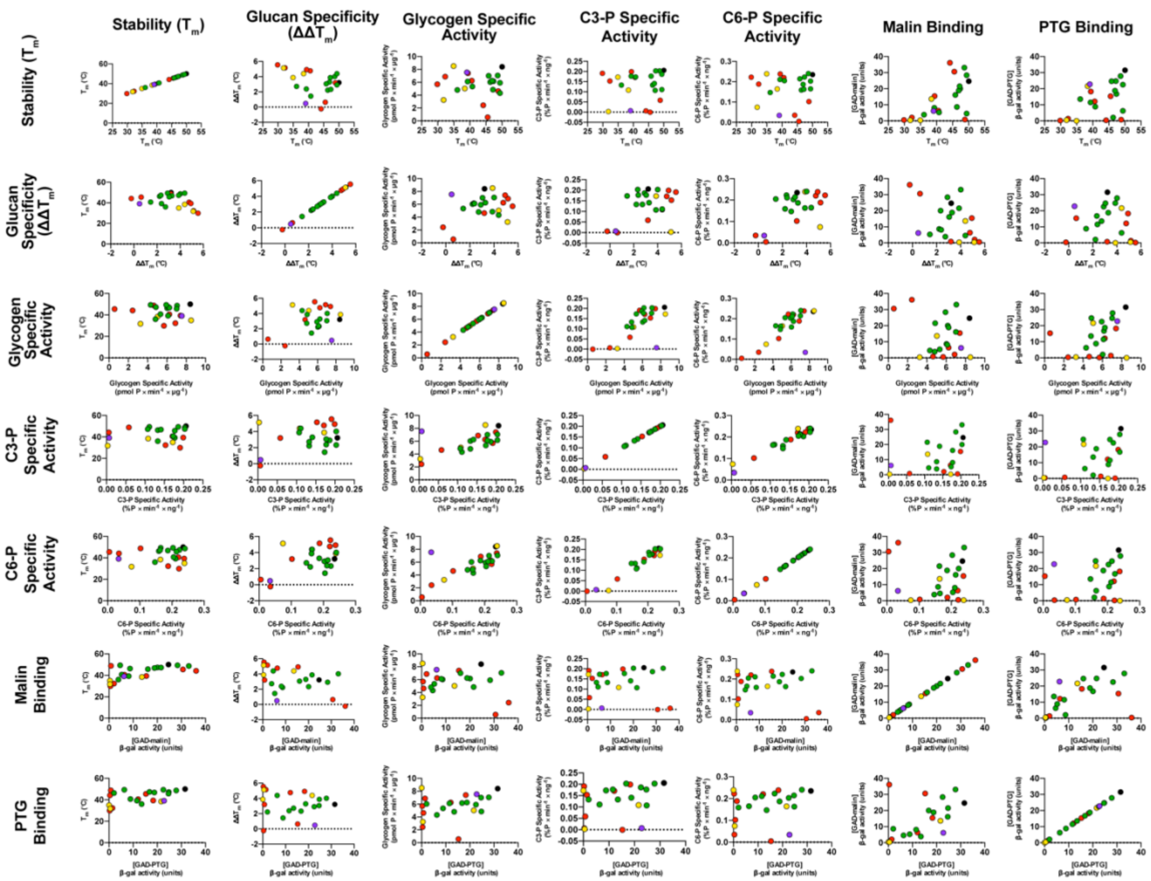


**Fig. S3. Pairwise correlation plots of all laforin functional measurements.** Nonparametric spearman correlations were used to compare laforin functions. R and p-values are in Table S1. Datapoints are color coded by structural group: WT (black); CBM (red); CBM-DSP interface (yellow); DSP (green); dimer interface (purple).

| | | **Stability ($T_m$)** | **Glucan specificity ($\Delta\Delta T_m$)** | **Glycogen specific activity** | **C3-P specific activity** | **C6-P specific activity** | **malin binding (β-gal activity)** | **PTG binding (β-gal activity)** |
|---|---|---|---|---|---|---|---|---|
| **Stability ($T_m$)** | Spearman r | 1 | -0.2955 | -0.005929 | 0.2141 | 0.09289 | 0.6087 | 0.499 |
| | P (two-tailed) | | 0.1711 | 0.9786 | 0.3266 | 0.6734 | 0.0021 | 0.0154 |
| | P value summary | | ns | ns | ns | ns | ** | * |
| **Glucan specificty ($\Delta\Delta T_m$)** | Spearman r | -0.2955 | 1 | 0.165 | 0.2195 | 0.414 | -0.417 | -0.1433 |
| | P (two-tailed) | 0.1711 | | 0.4518 | 0.3142 | 0.0495 | 0.0478 | 0.5143 |
| | P value summary | ns | | ns | ns | * | * | ns |
| **Glycogen specific activity** | Spearman r | -0.005929 | 0.165 | 1 | 0.6507 | 0.6897 | 0.01087 | 0.3646 |
| | P (two-tailed) | 0.9786 | 0.4518 | | 0.0008 | 0.0003 | 0.9607 | 0.0872 |
| | P value summary | ns | ns | | *** | *** | ns | ns |
| **C3-P specific activity** | Spearman r | 0.2141 | 0.2195 | 0.6507 | 1 | 0.9177 | 0.1637 | 0.3051 |
| | P (two-tailed) | 0.3266 | 0.3142 | 0.0008 | | < 0.0001 | 0.4556 | 0.1569 |

| | | | | | | | | |
|---|---|---|---|---|---|---|---|---|
| | P value summary | ns | ns | *** | | **** | ns | ns |
| **C6-P specific activity** | Spearman r | 0.09289 | 0.414 | 0.6897 | 0.9177 | 1 | 0.03656 | 0.1789 |
| | P (two-tailed) | 0.6734 | 0.0495 | 0.0003 | < 0.0001 | | 0.8685 | 0.4142 |
| | P value summary | ns | * | *** | **** | | ns | ns |
| **malin binding (β-gal activity)** | Spearman r | 0.6087 | -0.417 | 0.01087 | 0.1637 | 0.03656 | 1 | 0.6462 |
| | P (two-tailed) | 0.0021 | 0.0478 | 0.9607 | 0.4556 | 0.8685 | | 0.0009 |
| | P value summary | ** | * | ns | ns | ns | | *** |
| **PTG binding (β-gal activity)** | Spearman r | 0.499 | -0.1433 | 0.3646 | 0.3051 | 0.1789 | 0.6462 | 1 |
| | P (two-tailed) | 0.0154 | 0.5143 | 0.0872 | 0.1569 | 0.4142 | 0.0009 | |
| | P value summary | * | ns | ns | ns | ns | *** | |

**Table S1. Statistical results from pairwise Spearman correlation analysis.** Analysis were performed using Prism Graphpad software.
