## Supplemental File 1 for "An empirical pipeline for personalized diagnosis of Lafora disease mutations"

#### Ribbon Map of Laforin WT (in D%, Deuteration Level)

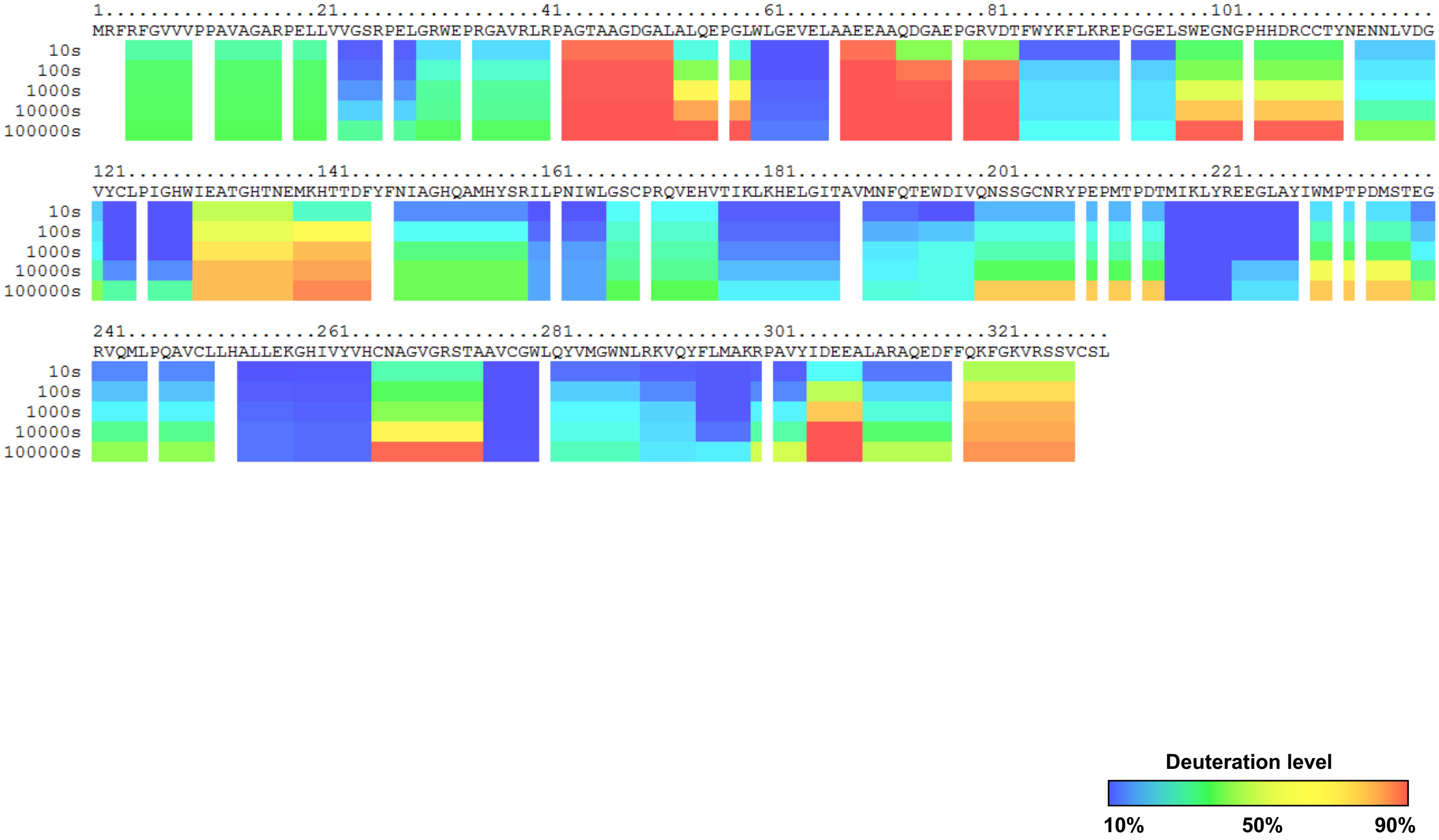

### Ribbon Map of Laforin R91P (in D%, Deuteration Level)

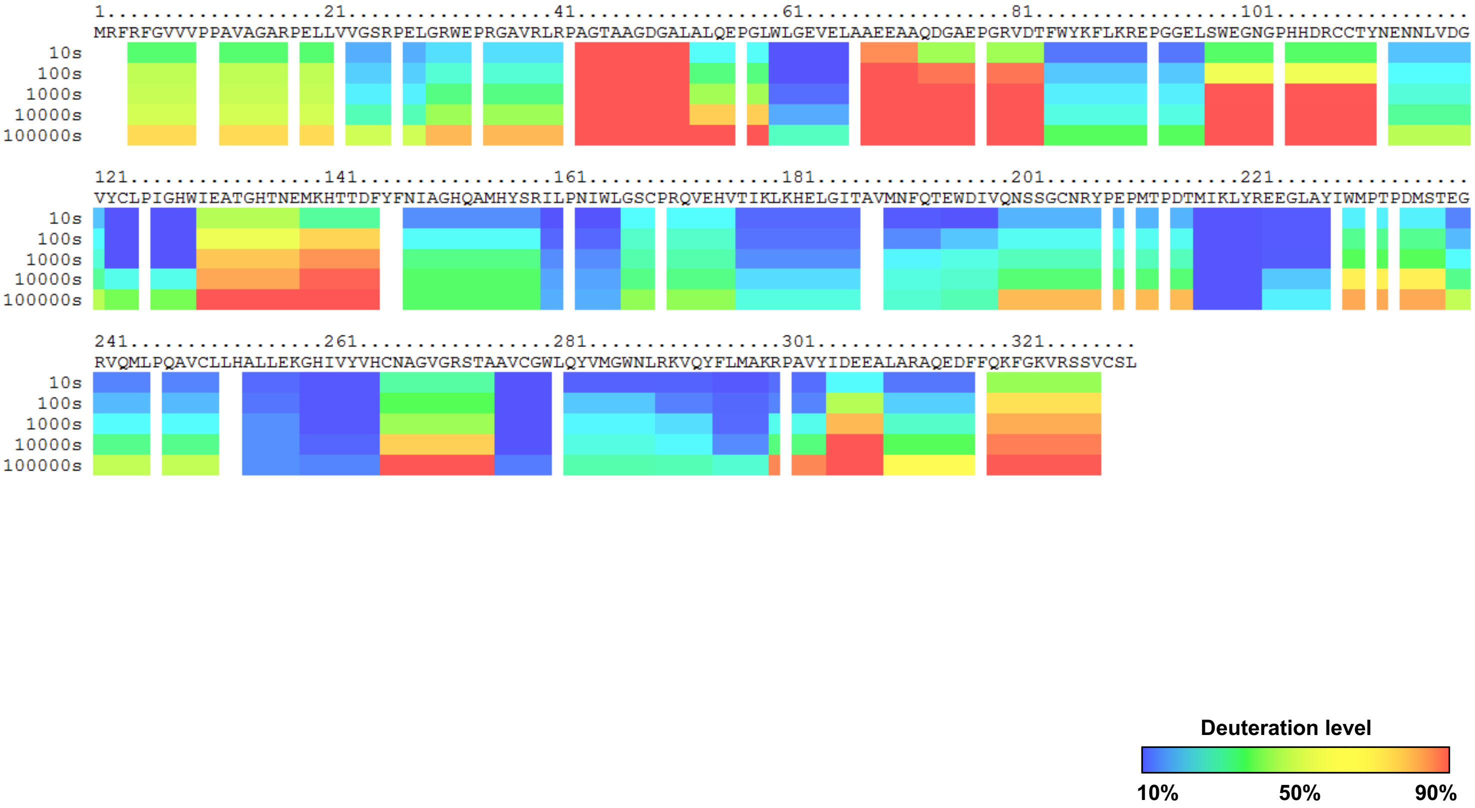

### Ribbon Map of Laforin P211L (in D%, Deuteration Level)

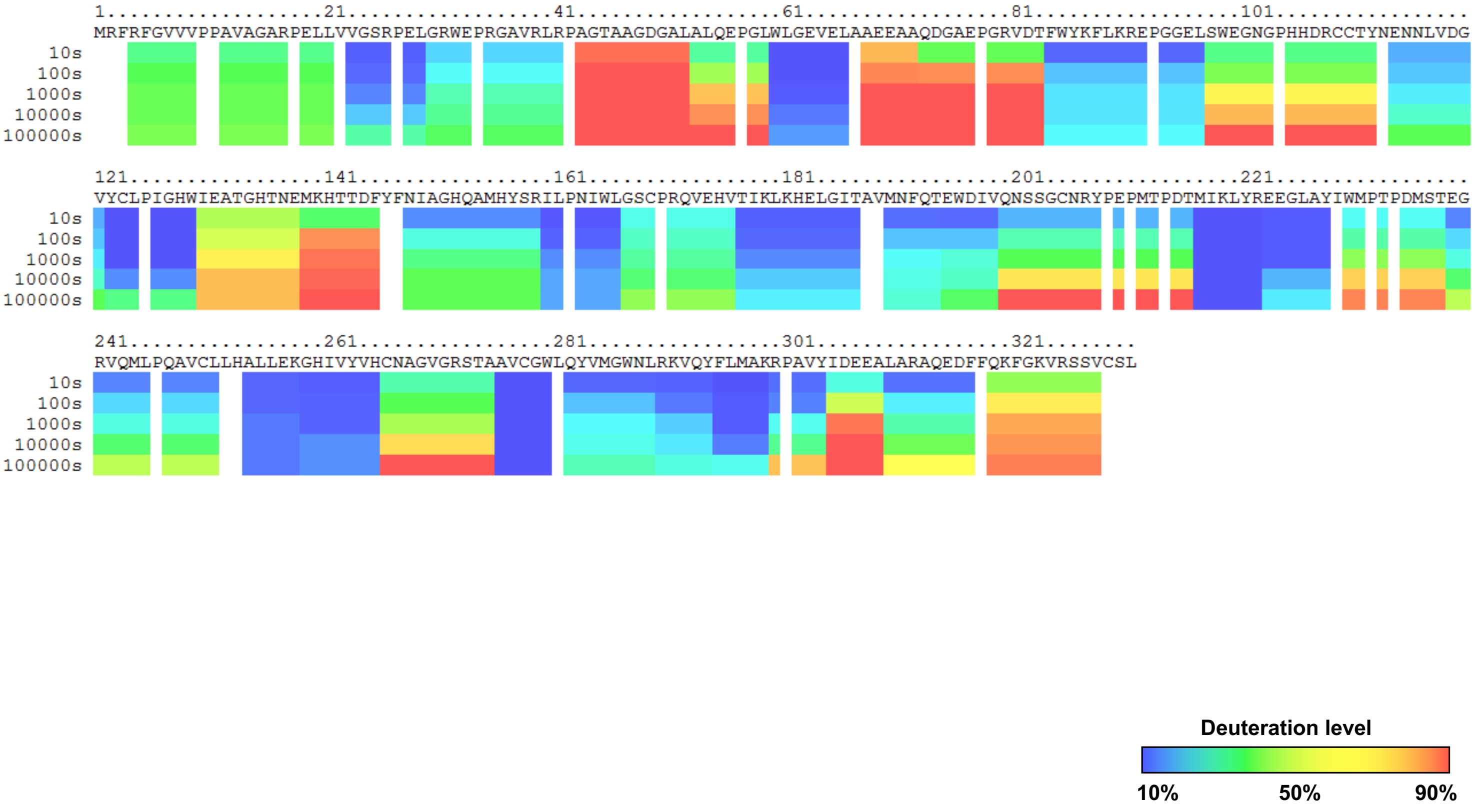

### Ribbon Map of Laforin G279C (in D%, Deuteration Level)

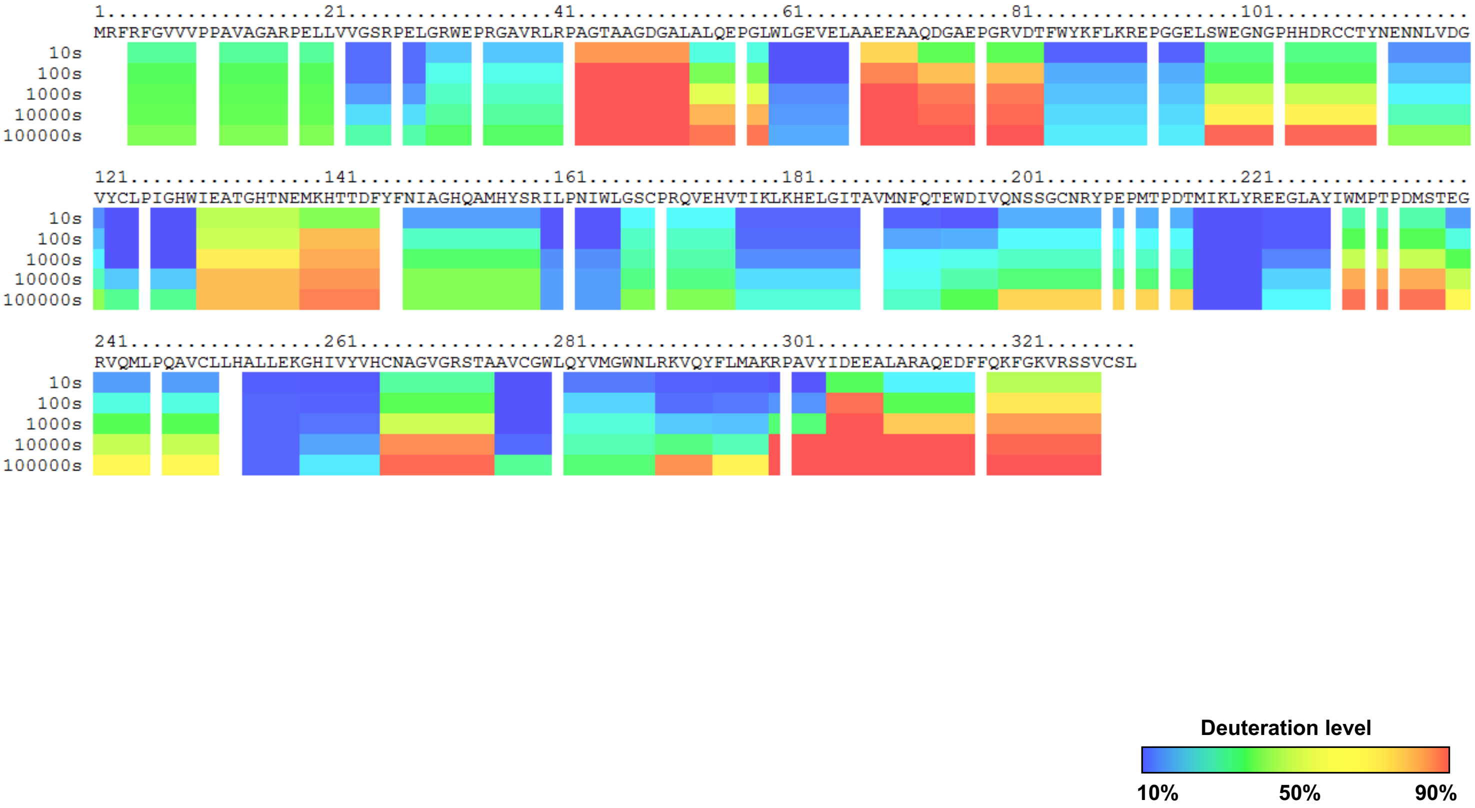

### Ribbon Map of Laforin F321C (in D%, Deuteration Level)

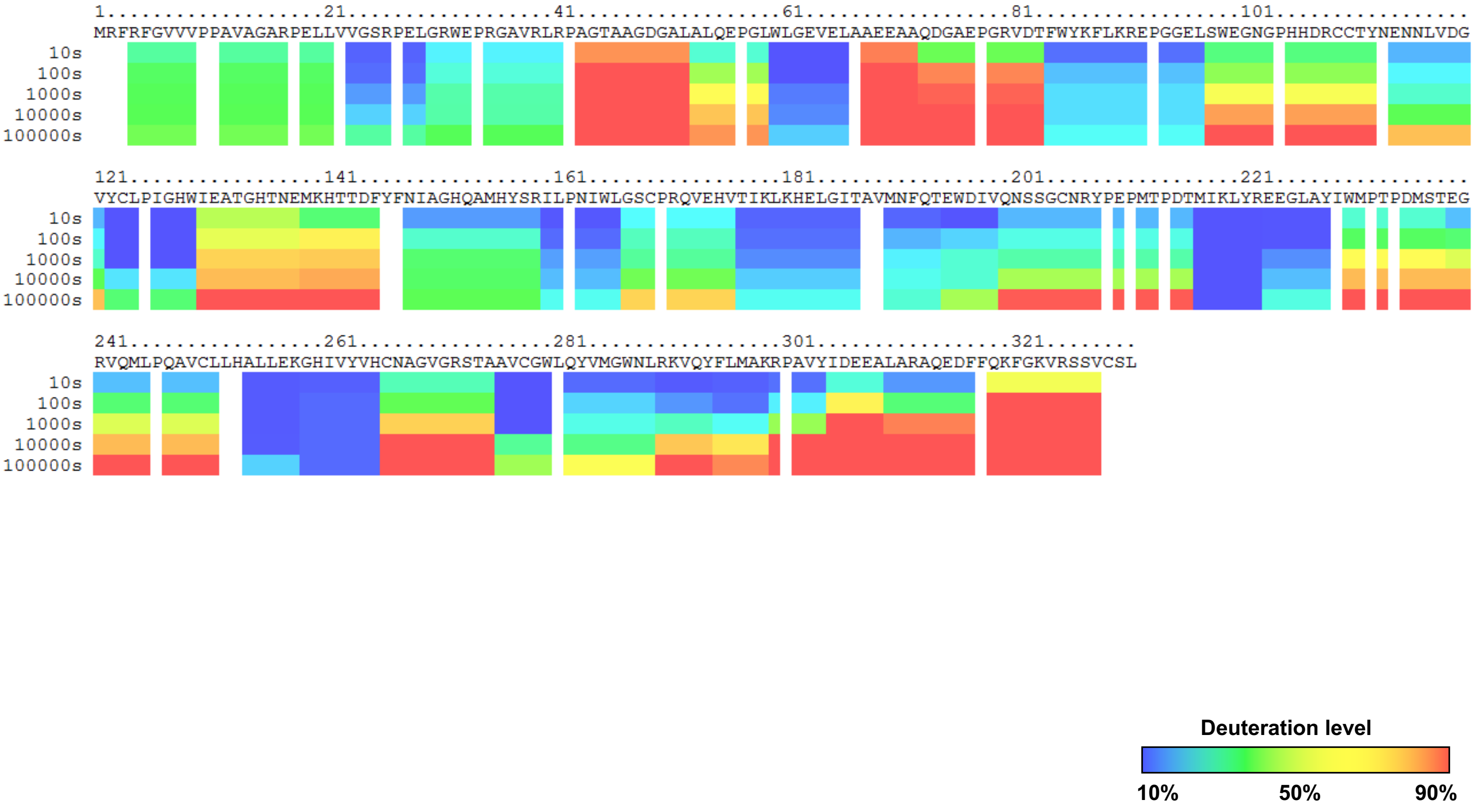
