## Supplemental File 2 for "An empirical pipeline for personalized diagnosis of Lafora disease mutations"

Difference Map (R91P- WT)(in D%, Deuteration Level)

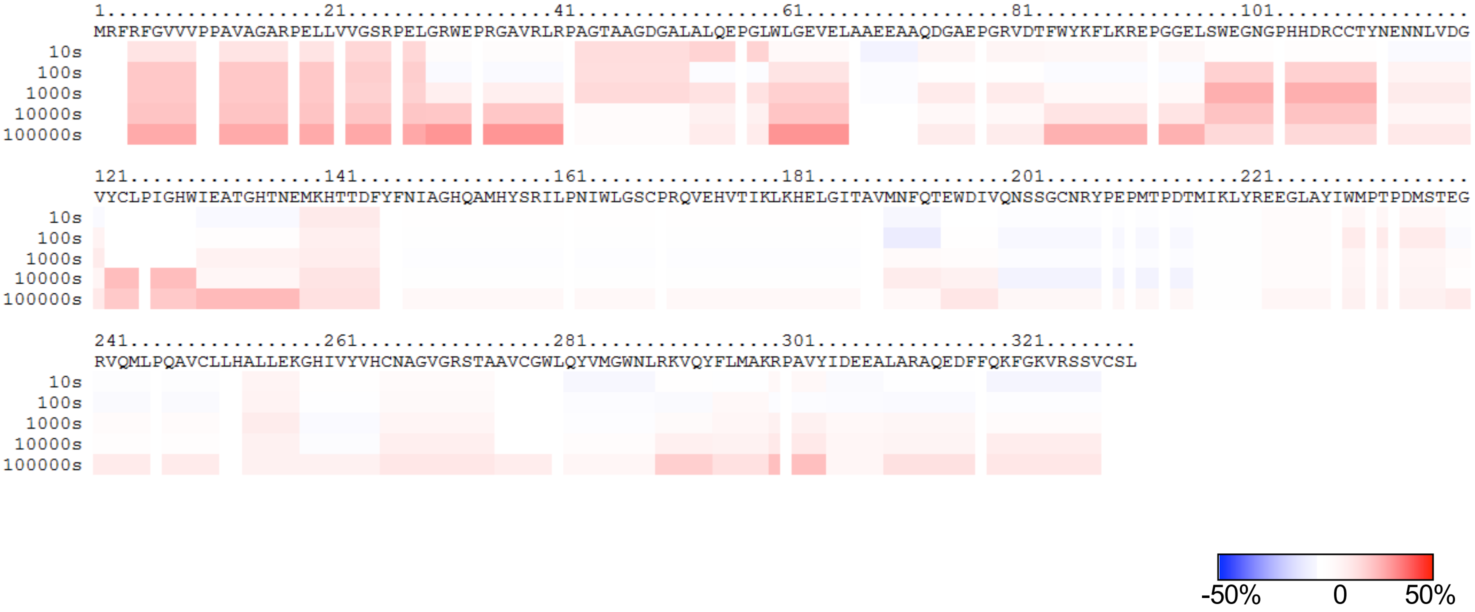


Blue indicates the regions which exchange slower in the mutant; red indicates the regions which exchange faster in the mutant.

Difference Map (P211L- WT)(in D%, Deuteration Level)

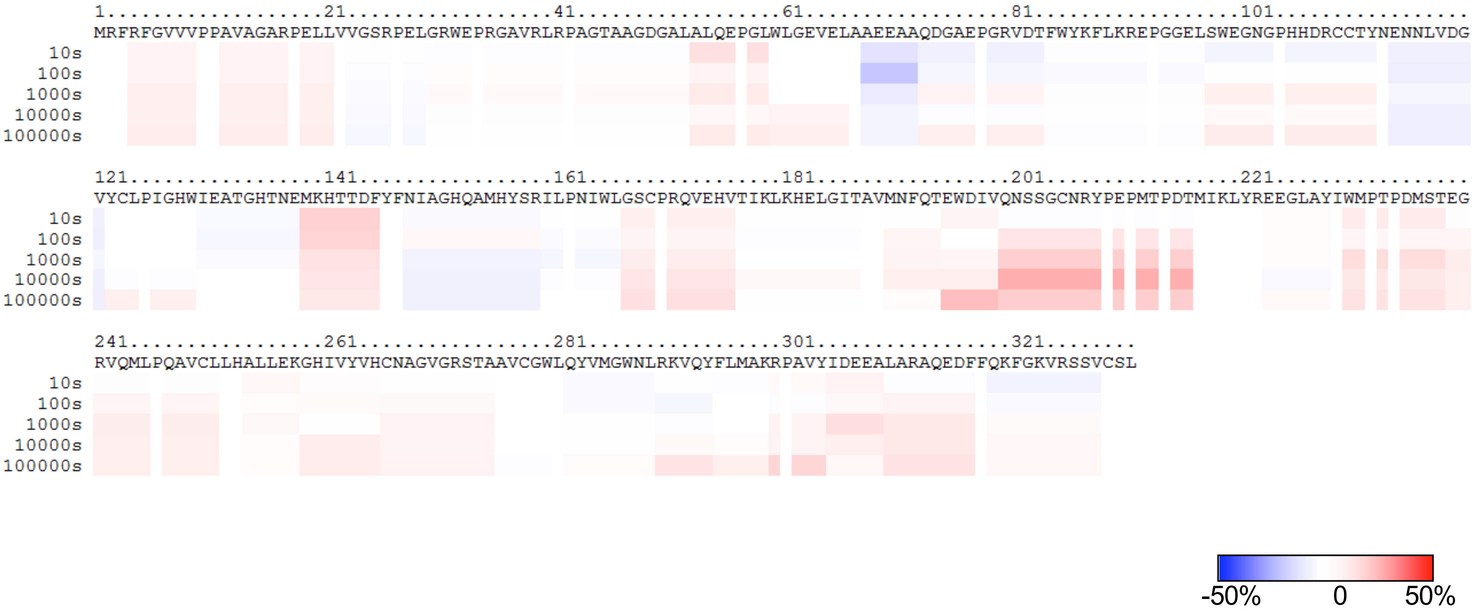


Blue indicates the regions which exchange slower in the mutant; red indicates the regions which exchange faster in the mutant.

Difference Map (G279C- WT)(in D%, Deuteration Level)

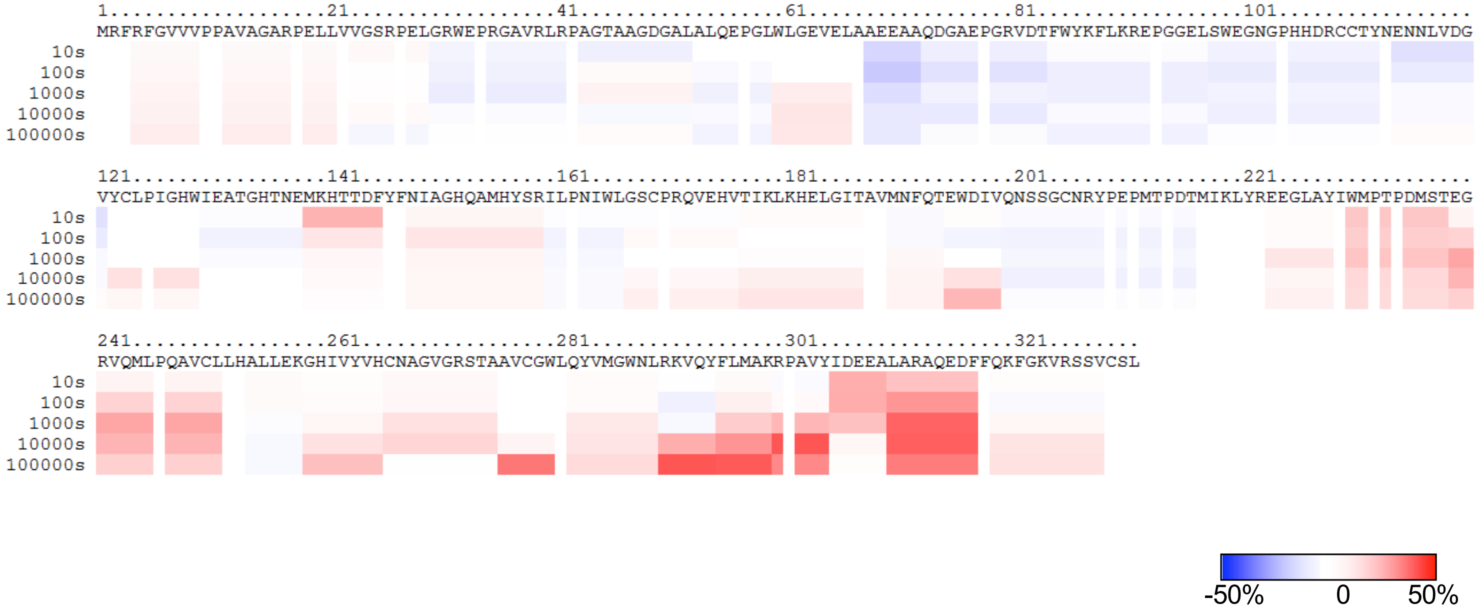


Blue indicates the regions which exchange slower in the mutant; red indicates the regions which exchange faster in the mutant.

Difference Map (F321C- WT)(in D%, Deuteration Level)

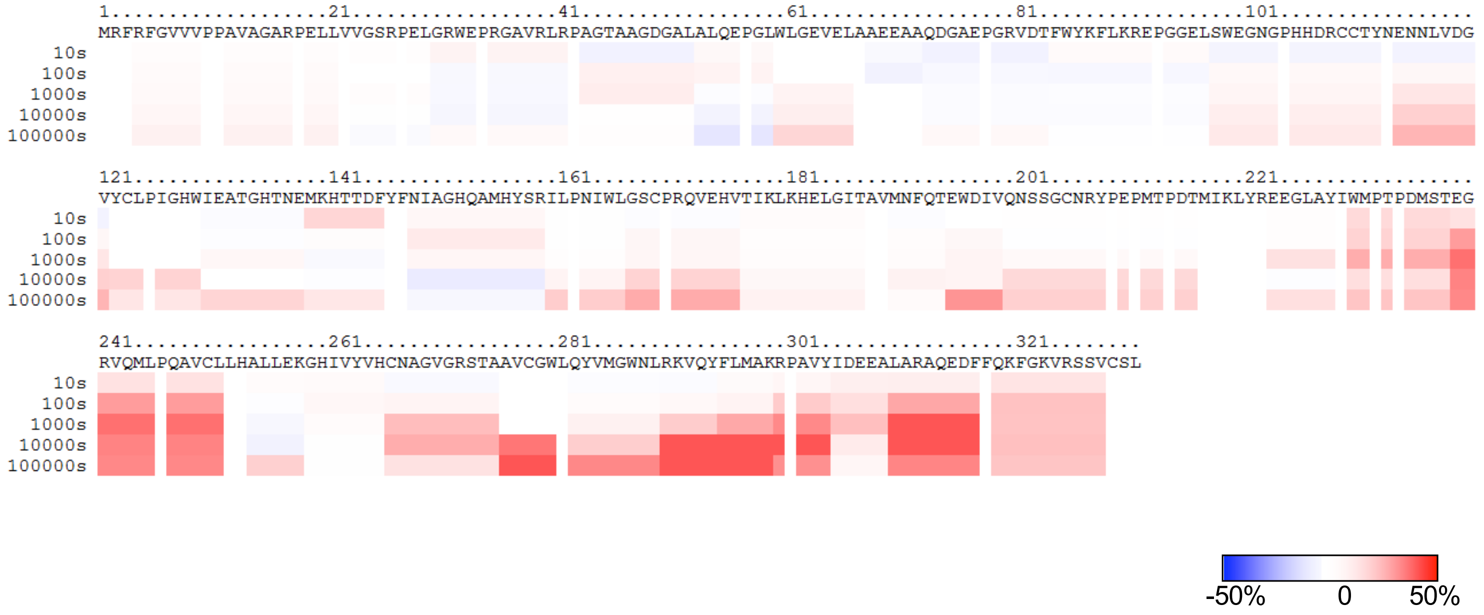


Blue indicates the regions which exchange slower in the mutant; red indicates the regions which exchange faster in the mutant.
